# Fungal communities living within leaves of native Hawaiian dicots are structured by landscape-scale variables as well as by host plants

**DOI:** 10.1101/640029

**Authors:** JL Darcy, SOI Switf, GM Cobian, G Zahn, BA Perry, AS Amend

## Abstract

A phylogenetically diverse array of fungi live within healthy leaf tissue of dicotyledonous plants. Many studies have examined these endophytes within a single plant species and/or at small spatial scales, but landscape-scale variables that determine their community composition are not well understood, either across geographic space, across climatic conditions, or in the context of host plant phylogeny. Here, we evaluate the contributions of these variables to endophyte beta diversity using a survey of foliar endophytic fungi in native Hawaiian dicots sampled across the Hawaiian archipelago. We used Illumina technology to sequence fungal ITS1 amplicons to characterize foliar endophyte communities across five islands and 80 host plant genera. We found that communities of foliar endophytic fungi showed strong geographic structuring between distances of seven and 36 km. Endophyte community structure was most strongly associated with host plant phylogeny and evapotranspiration, and was also significantly associated with NDVI, elevation, and solar radiation. Additionally, our bipartite network analysis revealed that the five islands we sampled each harbored significantly specialized endophyte communities. These results demonstrate how the interaction of factors at large and small spatial and phylogenetic scales shape fungal symbiont communities.

## INTRODUCTION

Foliar endophytic fungi (FEF), defined here as all fungi living within leaf tissue but not causing any outward signs of disease (sensu (Stone, Bacon, & White, 2000)) are effectively invisible and represent a “hotspot” of undescribed fungal diversity (Arnold & Lutzoni, 2007; Porras-Alfaro & Bayman, 2011). Indeed, a large portion of the vast undescribed fungal diversity on earth (Blackwell, 2011) are presumed to live cryptic lifestyles in association with plant hosts (Hawksworth & Rossman, 1997), particularly in the tropics where plant diversity is highest. Due to their cryptic lifestyles and high species richness (Blackwell, Hibbett, Taylor, & Spatafora, 2006; Rodriguez, White, Arnold, & Redman, 2009), many questions remain about which factors determine how FEF are distributed throughout nature.

Although all available evidence suggests that most eudicot FEF are horizontally transmitted and not inherited via seed (Bayman, Angulo-Sandoval, Báez-Ortiz, & Lodge, 1998), it is unclear which factors structure FEF community composition, and how those factors rank in their relative importance. Previous studies that have examined foliar fungal communities in this ecological and biogeographic context have noted several different drivers of community composition and biogeography. Temperature (Coince et al., 2014), geographic distance (U’Ren, Lutzoni, Miadlikowska, Laetsch, & Arnold, 2012), elevation and rainfall (Zimmerman & Vitousek, 2012), and vegetation density/urbanization (Massimo et al., 2015; Unterseher, Petzold, & Schnittler, 2012; Vincent, Weiblen, & May, 2016) are all putatively important variables for FEF community composition. We might attribute this diversity of results to idiosyncrasies of experimental design in which specific variables or host taxa were held constant to address focused hypotheses, and to the simultaneous influence of many possible factors that may not all be analyzed in any given study.

In fact, the bulk of FEF community research is represented by studies focusing on one or two specific host plants (Felber et al., 2016; González-Teuber, 2016; Kato, Fukasawa, & Seiwa, 2017; Mejía et al., 2014; Oono, Lefèvre, Simha, & Lutzoni, 2015; Polonio et al., 2015; Saucedo-García, Anaya, Espinosa-García, & González, 2014; Zimmerman & Vitousek, 2012), often to address hypotheses about how distance or climate impacts composition. Still, studies that have surveyed FEF or other leaf-associated fungi among multiple plant species have found that host identity strongly determines fungal composition (Huang, Devan, U’Ren, Furr, & Arnold, 2016; Kembel & Mueller, 2014; Massimo et al., 2015; U’Ren & Arnold, 2016; Unterseher et al., 2012; Vincent et al., 2016). Despite examples of studies at either broad phylogenetic scales or broad spatial scales, comparatively few studies combine these into a single analytical framework (but see (U’Ren et al., 2012, 2019)). By simultaneously examining a wide diversity of hosts over a wide diversity of habitats at the landscape scale, we might explicitly address how host identity interacts with the abiotic environment to structure FEF community composition, and the extent to which FEF are geographically restricted either by distance or by island barriers.

Here, we use the breadth of environmental and evolutionary diversity in the Hawaiian Archipelago to determine which factors structure FEF beta diversity in native Hawaiian dicots. Specifically, we test the hypotheses that elevation, rainfall, and host plant phylogeny are the strongest predictors of FEF community composition of native plants across Hawaii (Figure 1). We include additional potential explanatory variables in our analysis such as evapotranspiration (Kivlin, Hawkes, & Treseder, 2011; Lewis, Ravel, Naffaa, Astier, & Charmet, 1997) and geographic distance (Higgins, Arnold, Coley, & Kursar, 2014), known to be important determinants in other fungal systems. We found that FEF community structure was most strongly associated with host plant phylogeny and evapotranspiration, and was also significantly associated with NDVI, elevation, and solar radiation. Spatial structuring of FEF communities was strongest at distances between seven and 36 km.

**Figure 1:**
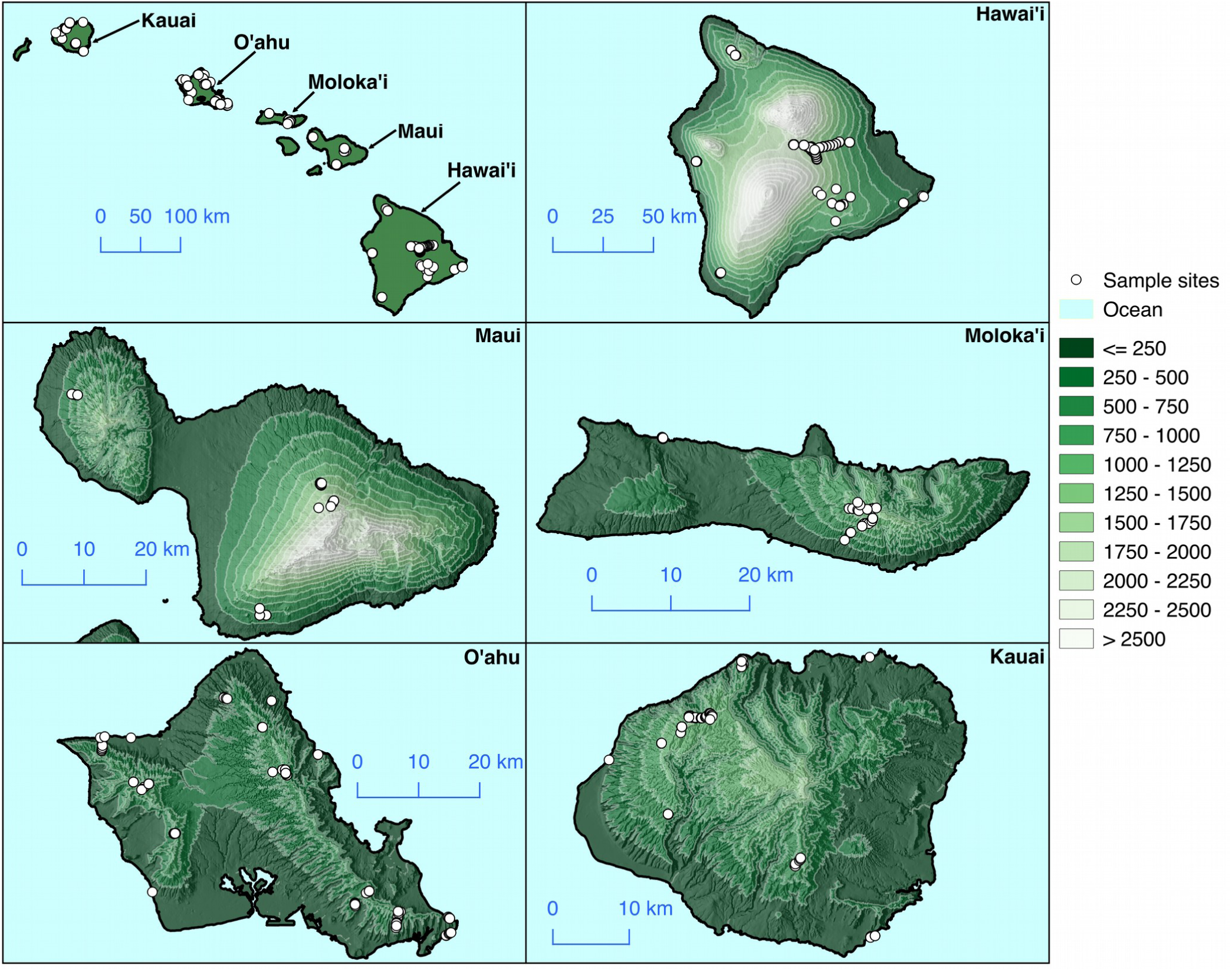
Map of sample locations across the Hawaiian Archipelago. Samples were collected from the five major islands in the Hawaiian archipelago: Hawai‘i, Maui, Moloka‘i, O‘ahu, and Kaua‘i. Sampling was most dense on Hawai‘i and on O‘ahu islands, where accessibility was easiest. Our sampling strategy was to use elevational transects where possible, in order to capture elevational and climatic variation. Elevations in figure legend are meters above sea level. This is visible in the transects on Hawai‘i, Moloka‘i, Kaua‘i, and O‘ahu, which run orthogonally to the topographic lines (white). Transects are less pronounced on Maui because of limited accessibility.

## MATERIALS AND METHODS

### Sample Collection

General sampling locations were selected to maximize habitat, phylogenetic and spatial diversity. Location selection prioritized access, a reasonable permitting process, and the known presence of ten or more native plant species. Sample collection took place between July 2014 and October 2016. Access was the most limiting factor in comparatively fewer sampling locations on Molokai and Maui Islands. We chose not to collect plants that are federally listed as threatened or endangered. At each location, the first occurrence of an apparently healthy, naturally recruited individual was selected for sampling. We only sampled native plants. Because some native species are comparatively common, and others rare, we limited our sampling to a single individual from each naturally occurring non-listed dicot at each location (*e*.*g*. a trail or a park), because we did not wish to confound our data with uneven spatial heterogeneity among sub-sampled plants. In most cases more than a single individual was collected per island since it occurred at more than one location. Numerous mature sun leaves were collected such that when combined, they covered a surface area roughly equivalent to two adult-sized hands. Life form and stature differed markedly across our sampled plants, so it was difficult to precisely standardize the height at which leaves were collected. When possible, leaves from trees and shrubs were collected at eye level and from at least four aspects of the tree canopy. We targeted dicots because they have horizontally transmitted FEF (compared to many vertically transmitted FEF in some monocots). Although not all host plant genera were collected at every sample location, the three most common genera we collected (*Metrosideros*, n=119; *Leptecophylla*, n=89; and *Vaccinium, n=*147) were each collected across elevations ranging from sea level to over 2000 meters above sea level. The location of each plant was recorded with a GPS and plants were positively identified in the field and/or vouchered for subsequent identification (vouchers deposited at Joseph F. Rock Herbarium at the University of Hawaii, Manoa; HAW). Leaves were refrigerated until subsequent processing (within 72 hours of collection). A total of 1099 samples were collected in this way, although not all were used in the analyses presented here (see results section).

We surface sterilized leaves to exclude fungi present on leaf surfaces. After rinsing in water, forty leaf discs were extracted per individual host by punching leaves with a sterile standard paper single hole punch (approximately 0.5 cm diameter). Hole punches were sterilized by washing with 70% ethanol and flamed between uses. For plants with very small leaves, entire leaves were used such that the area of those leaves was the same as the area of leaf discs. Leaf discs (or aforementioned collections of small leaves) were then placed into loose-leaf tea bags that were subsequently stapled shut, submerged in 1% NaOCl for two minutes, then 70% EtOH for two minutes, followed by two rinses with sterile water for two minutes each. Rinse water was included in extraction controls to verify sterility of surface water.

### DNA Isolation

Ten leaf discs per DNA extraction were placed in MP Biomedical Lysing Matrix A tubes (MP Biomedical, Santa Ana, CA, USA) containing DNA isolation solutions from the Mo Bio PowerPlant Pro DNA Isolation kit (Solution PD1, Solution PD2, Phenolic Separation Solution, and RNase A Solution; Mo Bio, Carlsbad, CA, USA, a subsidiary of QIAGEN). Leaf discs were homogenized using a Mini-Beadbeater 24 (BioSpecs Inc. OK) at 3,000 oscillations per min for two minutes. Lysate was centrifuged at 13,000 RPM for two minutes and transferred to individual wells of a MoBio PowerPlant Pro DNA 96-well Isolation kit for subsequent extraction following the manufacturer’s protocol.

### PCR Amplification and Illumina Library Preparation

We amplified the ITS1 region of the ribosomal cistron using fungal specific primers ITS1F (5’-CTTGGTCATTTAGAGGAAGTAA-3’) and ITS2 (5’-GCTGCGTTCTTCATCGATGC-3’), along with Illumina adapters and Golay barcodes incorporating dual indexing, using previously published thermal cycling parameters (Smith & Peay, 2014). PCRs were carried out in 25 μl reactions using the KAPA3G Plant PCR kit (KAPAl reactions using the KAPA3G Plant PCR kit (KAPA Biosystems, Wilmington, MA, USA), nine ul of DNA extract (concentration not measured) and 0.2 μl reactions using the KAPA3G Plant PCR kit (KAPA M each of the forward and reverse primers. Negative PCR and extraction controls were included. PCR products were purified and normalized using just-a-plate 96 PCR Purification and Normalization Kit (Charm Biotech, San Diego, California, USA). Normalized PCR products were pooled and concentrated using a streptavidin magnetic bead solution. Pooled PCR products were sequenced on three separate reactions in order of processing completion, using the 2×300 paired-end (PE) sequencing protocol on an Illumina MiSeq sequencing platform (Illumina Inc., Dan Diego, CA, USA). We did not include controls for sequencing run variation, and we acknowledge this as a limitation of our study. We also did not quantify DNA concentration of extracts, but the amplification-normilization experimental design allows for characterization of microbial communities even from inefficient DNA extractions. We do not think this is a significant source of bias in our analysis, since negative controls (which contained too little DNA to amplify) produced a paucity of sequence data when compared to samples, almost all of which produced many thousands of sequence reads. Still, it is possible that variation in extraction efficiency could result in biased alpha diversity estimates. Due to low sequencing success, negative controls were not accounted for in sample processing.

### DNA Sequence data processing and bioinformatics

QIIME v1.9.1 (Caporaso et al., 2010) was used to demultiplex raw DNA sequence data into individual fastq files for each sample. Although paired-end sequencing was used, only the R1 read (corresponding to primer ITS1f) was used for downstream analysis, since sequencing quality of reverse reads was generally poor after the first 150 nucleotides. VSEARCH v2.14.2 (Rognes, Flouri, Nichols, Quince, & Mahé, 2016) was used to discard reads with an average quality score below 25 (Illumina Q+33 format), then ITSx v1.1 (Bengtsson-Palme et al., 2013) was used to extract the ITS1 region from quality-filtered files. Sequence data were filtered by average score rather than maximum score, because nucleotides with poor quality scores that were not within the ITS1 region could disqualify sequences that would be otherwise high-quality within the ITS1 region. In order to eliminate variability in sequences caused by sequencing and PCR error, ITSx-processed sequences were de-noised using DADA2 v1.12 (Callahan et al., 2016) using the authors’ recommended settings (maximum expected errors = 2, maximum Ns = 0). Separate DADA2 error rates models were used for each sequencing run in our data set, although the models appeared qualitatively similar. De-noised sequences were then clustered into 95% OTUs using VSEARCH (Rognes et al., 2016).

Taxonomy was assigned to each OTU using the UNITE database (v7) (Nilsson, 2011) and QIIME’s assign_taxonomy.py script (Caporaso et al., 2010) with the BLAST method using the default maximum e-value of 0.001. OTUs not belonging to kingdom Fungi were discarded. Our OTU table was then rarefied (*i*.*e*. randomly downsampled) to 1500 sequences per sample, discarding samples with fewer than 1500 sequences or samples for which host plants could not be satisfactorily identified. We chose the depth of 1500 sequences as a compromise between a depth at which almost all samples had plateaued on our collector’s curves (Supplementary figure S1), and a level that resulted in the fewest discarded samples. Filtering and rarefaction were performed in R v3.5.0 (R Core Team, 2014) using custom software written by the authors.

Ghost Tree v2019 (Fouquier et al., 2016) was used to construct a phylogenetic tree for the remaining OTUs. Briefly, Ghost Tree allows phylogenetic trees to be made from ITS1 sequence data, which are often un-alignable at high taxonomic ranks. This is done using a backbone tree created with the 18S rRNA gene, and then ITS1 sequences are used to refine the tree at a phylogenetic scale where those sequences can be meaningfully aligned (*e*.*g*. genus level). In an analysis of real and simulated ITS1 data, Fouquier et al. (2016) found that using UniFrac distances calculated using a tree made with Ghost Tree were better explained by environmental variables in ANOSIM models than were distances calculated using phylogeny-independent metrics. (Fouquier et al., 2016). This method was also successfully used to study differences in fungal community composition in the human infant microbiome (Ward et al., 2018) and in a study of FEF in grass as well (Lumibao et al., 2019). We used a Ghost Tree that was made using the SILVA database (v132) for the 18S backbone, and the UNITE database (v8). was used for ITS refinement. Tips of the Ghost Tree were renamed with OTU identifiers where OTUs were assigned taxonomy to a UNITE entry in the Ghost Tree. In cases where multiple OTUs were assigned to the same UNITE entry, a polytomy was created to fit those OTUs into the tree. This method may exclude novel taxa that are not present in the UNITE reference database, and this is a limitation of all studies using such a database, including this one.

The Ghost Tree was used with weighted UniFrac (Lozupone & Knight, 2005) (hereafter referred to as “UniFrac”) to construct a beta diversity distance matrix for the samples. UniFrac was used because it is downweights additional branch length contributed by slight intragenomic variation among tandem repeats. Even if an individual putative fungal species contains several OTUs, each will only contribute a negligible amount of branch length to a sample, as compared to non-phylogenetic metrics (*e*.*g*. Bray-Curtis), which would consider those OTUs as different as any other pair. Because UniFrac community dissimilarity considers the shared phylogenetic branch-lengths between two communities, it is robust to the case where an OTU is only found within one sample, and is similarly robust to the case where samples do not share any OTUs, which can be problematic for other community dissimilarity metrics where overall diversity is much higher than within sample diversity. This is important for our analysis of FEF communities, because many previous studies have shown that FEF are hyperdiverse even at local scales (*e*.*g*. within one host or within one hundred meters) (Jumpponen & Jones, 2009, 2010; Zimmerman & Vitousek, 2012). We suspected that this large amount of diversity would result in many pairs of samples that shared few or zero OTUs, which would result in an inflation of 1-values (maximum dissimilarity) when using non-phylogenetic beta diversity metrics such as Bray-Curtis or Jaccard community dissimilarity. Indeed, this same issue has been discussed by previous users of Generalized Dissimilarity Modeling (GDM) as a saturation of maximal dissimilarities, and was resolved by using phylogenetic dissimilarty metrics (Warren, Cardillo, Rosauer, & Bolnick, 2014). We confirmed that this was indeed the case, and that UniFrac distance values were normally distributed but Bray-Curtis dissimilarities were severely 1-inflated (Supplementary figure S2). For this reason, we used the UniFrac distance matrix for the remainder of our analysis.

### Spatiotemporal data

Using sample geographic coordinates and collection dates, environmental and climatic data for each sample were extracted from GIS layers using the R packages ‘*raster*’ v2.9 (Hijmans et al., 2014) and ‘*rgdal*’ v1.4 (Pebesma, Rowlingson, & Rogerbivandnhhno, 2012). Data were extracted from rasters generated for the same month that samples were collected (except for elevational data). Supplementary table S1 shows the sources of each GIS layer. Many explanatory variables were obtained from the Rainfall of Hawai‘i and Evapotranspiration of Hawai‘i websites (T W Giambelluca et al., 2014; Thomas W. Giambelluca et al., 2013), and elevation data were obtained using the USGS EarthExplorer online tool (http://earthexplorer.usgs.gov), courtesy of NASA EOSDIS Land Processes Distributed Active Archive Center and the United States Geological Survey’s Earth Resources Observation and Science Center. These explanatory variables were chosen either because previous studies of Hawaiian FEF had identified them as important (elevation, rainfall)(Zimmerman & Vitousek, 2012), because they were significant drivers of Hawaiian plant composition (slope, aspect), or because they made intuitive sense in the context of fungi that live within leaves (solar radiation, transpiration, evapotranspiration, leaf area index, NDVI). Slope and aspect of each sampling location were calculated from elevation raster data using the terrain function in ‘*raster*’. Data for aspect (the direction a sampling location faces) were converted into a distance matrix using the smallest arc-difference between any two given aspects. This was done because Euclidean distance is unsuitable for a measurement like aspect, where 355° is closer to 1° than it is to 340°. All variables are mean monthly values. NDVI (normalized difference vegetation index) is an index calculated from the amount of infra-red light reflected by plants, which is normalized using multiple wavelengths of visible light. This allows for discrimination between habitat types that are deferentially vegetated. Similarly, leaf area index is a measure of surface area of leaves (one-sided) per unit area of ground, and while it does not discriminate between different types of vegetation as does NDVI, unlike NDVI it measures the density of leaf surface area (habitat for FEF).

### Host plant phylogeny

A distance matrix of host plant phylogenetic distances was created using the angiosperm phylogeny of Qian and Jin (Qian & Jin, 2016). This distance matrix was made because the modeling method we use here (GDM, below) can accommodate distance matrices as explanatory variables, allowing for a phylogenetic distance matrix of hosts to be used instead of a simplified data structure such as a principal components vector or an array of categorical taxon identities. For each pair-wise comparison between two samples, pair-wise host plant phylogenetic distance was calculated as the mean cophenetic (branch-length) distance between members of the plant genera that were sampled, using the function cophenetic.phylo in the R package ‘*ape*’ v5.2 (Paradis & Schliep, 2019). In cases where host plant genus was not included in the phylogeny, the genus was substituted for the most closely-related genus available. Four genera were substituted in this way out of 80 total genera: *Labordia* → *Logania, Touchardia* → *Urtica, Waltheria* → *Hermannia, Nothocestrum* → *Withania*. We chose to resolve host plants at the genus level because species-level taxonomy can be unstable for recent adaptive radiations, and many Hawaiian species-level phylogenies are not resolved (Ziegler, 2002).

### Geospatial analyses

In order to test and visualize the spatial structuring of FEF beta diversity, we constructed a geographic semivariogram (Bachmaier & Backes, 2008) in R using a matrix of log_10_ transformed geographic distances between samples, and the UniFrac distance matrix of FEF community differences between samples. Semivariograms are useful tools in spatial ecology because they visualize the extent to which beta diversity exhibits spatial connectedness. They can show at what geographic range community composition is homogeneous, and at what range community composition is structured by distance. See Robeson et al. (2011) for a thorough discussion of the applicability of Semivariograms in this context, models that can be fit to them, and what hypotheses they can be used to test. Distance classes in the semivariogram were constructed using a sliding window methods, where each class shared two-thirds of its points with the previous class (Darcy, King, Gendron, & Schmidt, 2017). A logistic curve was fit to semivariance values and mean geographic distances within semivariogram distance classes using the Nelder-Mead algorithm, minimizing the sum of squared differences in beta diversity semivariance. The curve’s derivative was then used to find the geographic distance class that accounts for 99% of geographic distance where beta diversity is spatially autocorrelated. Since the derivative function of a logistic curve is a unimodal distribution, the 99% intervals of that function yield three distance classes for the logistic function: the first plateau, the class corresponding to 99% of the change in slope for the curve, and the second plateau. Mantel correlations were then performed on data within each of the three resulting distance classes.

Generalized dissimilarity modeling (GDM) (Ferrier, Manion, Elith, & Richardson, 2007) was used to model FEF beta diversity based on climatic factors (Supplementary table S1), geographic distance, host plant phylogeny, and DNA sequencing run. The goal of this analysis was to discover the relative importances of environmental variables on FEF community composition, and at what scales the variables explain FEF community composition (*e*.*g*. are small changes in a variable important, or only large changes). GDM is a form of non-linear matrix regression that is well-suited to statistical questions involving dissimilarity matrices (*e*.*g*. our UniFrac distance matrix, host plant cophenetic distance matrix, geographic distance matrix, and aspect arc-difference matrix). Unlike pair-wise Mantel tests or PerMANOVA, which make use of similar data, GDM can quantify the relative importance of environmental and geographic variables on community dissimilarity, even when the functional relationship between community dissimilarity and the environment is nonlinear (Fitzpatrick et al., 2013; Warren et al., 2014). Furthermore, GDM is effective because it models community dissimilarity associated with a given predictor variable while holding all other predictor variables constant (Fitzpatrick et al., 2013; Landesman, Nelson, & Fitzpatrick, 2014). For example, this enables GDM to model the effect of elevation while accounting for the effect of host plant phylogenetic distance. Because our data were generated using three separate DNA sequencing runs, we included “run” as a categorical variable in our model by creating a distance matrix where samples from the same run had distance=0, and samples of different runs had distance=1, as recommended by GDM’s creators (Ferrier et al., 2007).

We used backward elimination as implemented in the ‘*gdm*’ v1.3.11 R package to build a model, and then to simplify the model by removing minimally predictive variables. We began this process with a GDM model excluding variables that were correlated with another variable in the analysis at rho > 0.70 (Spearman). Leaf area index, wet canopy evaporation, temperature, cloud frequency, and relative humidity were removed this way (Supplementary figure S3). Log-transformed geographic distance between samples, host plant phylogenetic distance, sequencing run, and aspect arc-difference matrix were included as predictor matrices in the GDM, with FEF UniFrac distance as the response matrix. We then used GDM to test each variable within the model for significance using a permutation test. During this iterative process, the variable with the highest P-value was eliminated, and then the model was recalculated. This process was repeated until all remaining variables were statistically significant (*P* < 0.05). Sequencing run and transpiration were eliminated this way. The final GDM model was also analyzed using Bray-Curtis dissimilarities instead of UniFrac distances, to illustrate how the severe 1-inflation observed with Bray-Curtis was an impediment to model fitting.

Principal coordinate analysis (PCoA) was used to validate GDM results using an alternate statistical approach. PCoA vectors were calculated from the UniFrac distance matrix using the R package ‘*vegan*’ v2.5 (Oksanen et al., 2016). Principal Coordinates of Neighbor Matrices (PCNM)(Borcard & Legendre, 2002) was used on geographic and host phylogenetic distance matrices, for dimensionality reduction and to account for nonlinear features within those data. The same data used in our final GDM, except with geography and host represented by their first six PCNM vectors, were fit to FEF PCoA and tested using the ‘envfit’ function (Oksanen et al., 2016).

### FEF-island specialization analysis

Bipartite network analysis was used to test the extent to which each island (Figure 1) harbored specific FEF (FEF-island specialization). The d’ (“d prime”) statistic was calculated for each island using the OTU table using the ‘*bipartite*’ package v2.11 in R (Dormann, Gruber, & Fruend, 2008). d’ is a measure of network specialization that ranges from zero to one, where zero is perfect cosmopolitanism (all species are evenly shared among islands) and one is perfect specialization (each species is specific to only one island). d’ is calculated using a contingency matrix where each row is a unique lower-level group (island) and each column is a unique higher-level group (OTU), but in our OTU table each island contains multiple samples, so the number of observations per island is not consistent. To remedy this, we calculated d’ by aggregating all samples from the same island into one large sample (column sums), rarefied this aggregated table using the same depth to which samples were rarefied above (1500 observations), then calculated d’ values. Empirical d’ values were compared to null d’ values generated with the Vázquez null model, which is a fixed-connectance null model for bipartite networks (Vázquez et al., 2007), implemented as the vaznull function of ‘*bipartite*’. The procedure of aggregating, calculating an empirical d’, and calculating a Vázquez null d’ was repeated 1000 times in order to obtain bootstrapped distributions of empirical and null d’ values for each island. Statistical significance for d’ values for each island was tested using Welch’s unequal variance *t*-test to compare empirical and island null d’ values. This test was 2-tailed, since d’ could be significantly lower than the null distribution indicating cosmopolitanism, or significantly higher indicating specialization.

A similar procedure was applied to host plant genera in place of fungal OTUs (plant-island specialization). Since host is only observed once per sample, d’ was not bootstrapped. This was done in order to evaluate the extent to which our selection of host plants was significantly specialized to islands.

To evaluate whether FEF-island specialization could be attributed to island-specific plant selections, we also tested FEF-island specialization under the constraint of a generically identical plant community (at genus level) for each island. The same procedure for analyzing FEF-island specialization above was carried out, but during the aggregation step of each bootstrap, random samples for each island belonging to a fixed plant community were chosen to the exclusion of all others. This fixed community was defined by the plant identification overlap among all five islands. All d’ values calculated this way were calculated from the same plant community from each island. Analysis of these d’ values was calculated as above.

## RESULTS

### Sequence data

Samples were collected across a wide range of climatic conditions (Figure 2), representative of the distributions of those conditions in Hawai‘i. Our data set comprised 896 samples that passed quality-filtering and ITS1 extraction, consisting of 7002 OTUs. After non-fungal OTUs were discarded, samples were rarefied to 1500 sequences per sample, resulting in 3458 OTUs across 760 samples. Of those samples, 417 were from Hawai‘i, 84 from Kaua’i, 56 from Maui, 72 from Moloka‘i, and 131 from O‘ahu. Mean richness (number of OTUs observed) per sample was 27.3 with a standard deviation of 18.9 (Supplementary Figure S1). Geographic and botanical information for each sample used in our analysis is available as Supplementary Figure S4 and is available from our FigShare repository (see Data Accessibility section). Of 27 sequencing negative controls included in the study, 13 resulted in sequence data, with an average of 123 reads and 2.2 OTUs per sample. This sequencing depth was less than 1% of that found in the average biological sample (12,860). Because we were unable to trace the source of these contaminants (whether tag switching, cross contamination or reagent contamination) and because they amplified so poorly compared to biological samples, we did not attempt to adjust our data set to account for these. A data table containing geographic and host taxonomic information for each of the 760 samples used in our analysis is available as Supplementary Table S2.

**Figure 2:**
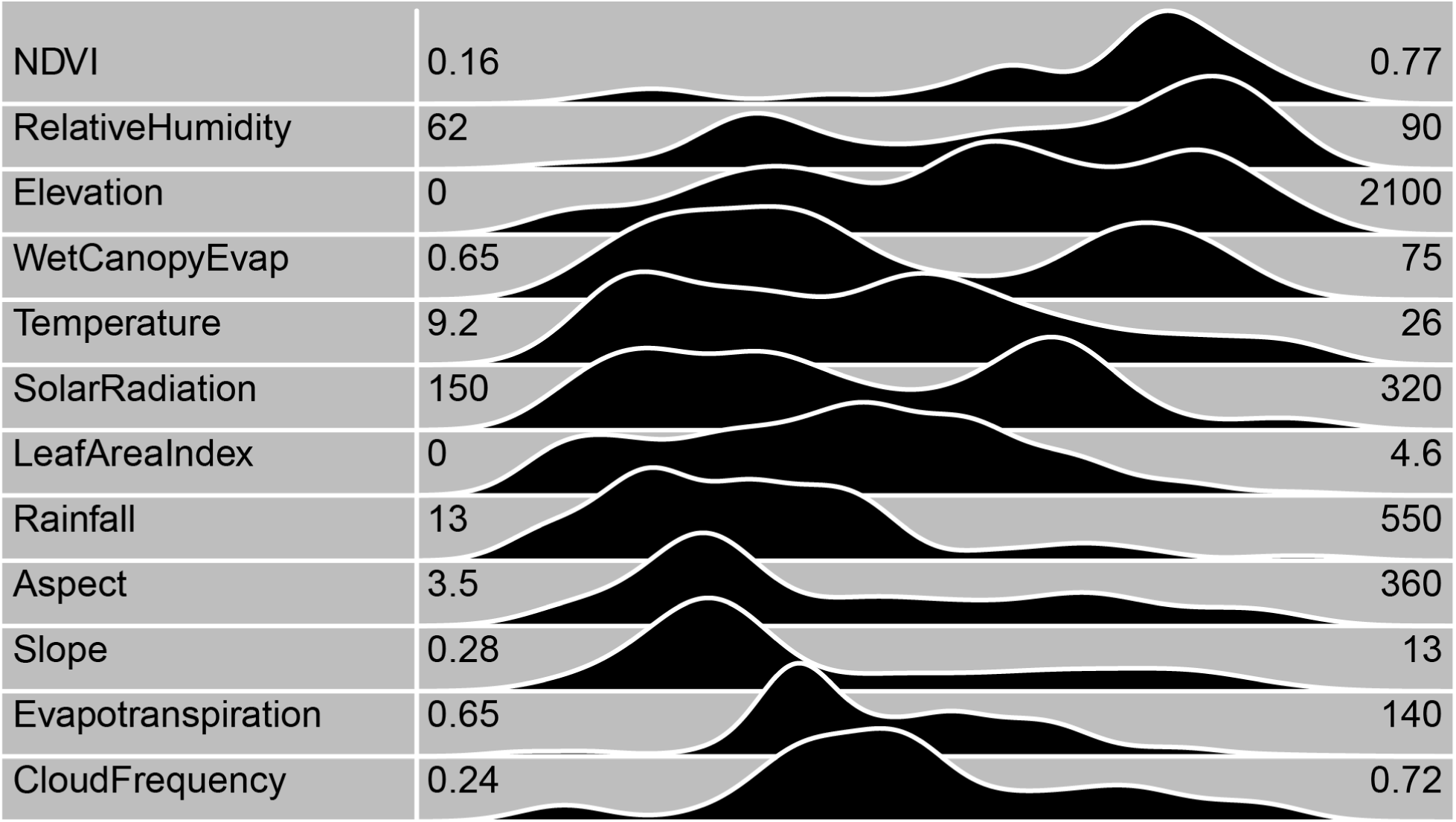
Ranges and distributions of explanatory variables. In this figure, each variable’s distribution across 722 samples is shown as a smoothed density curve between its range (numbers to left and right of curves, height of curve is relative frequency of observation). Units for range values are shown in Supplementary table S1. Each variable included in our analysis covered a wide range of environmental heterogeneity.

### Geospatial results

Our semivariogram analysis showed that FEF beta diversity is spatially structured between 7.6 and 36.3 km, where it reaches its autocorrelation distance (Figure 3). This plateau and subsequent noise in beta diversity semivariance occurs when FEF beta diversity is no longer spatially autocorrelated (Robeson et al., 2011). In our analysis, this occurs at geographic distances of 36.3 km. Within the distance class where beta diversity was spatially autocorrelated, *r*_*M*_ (Mantel correlation) was 0.16. In the other two distance classes, *r*_*M*_ was closer to zero as expected (Figure 3).

**Figure 3:**
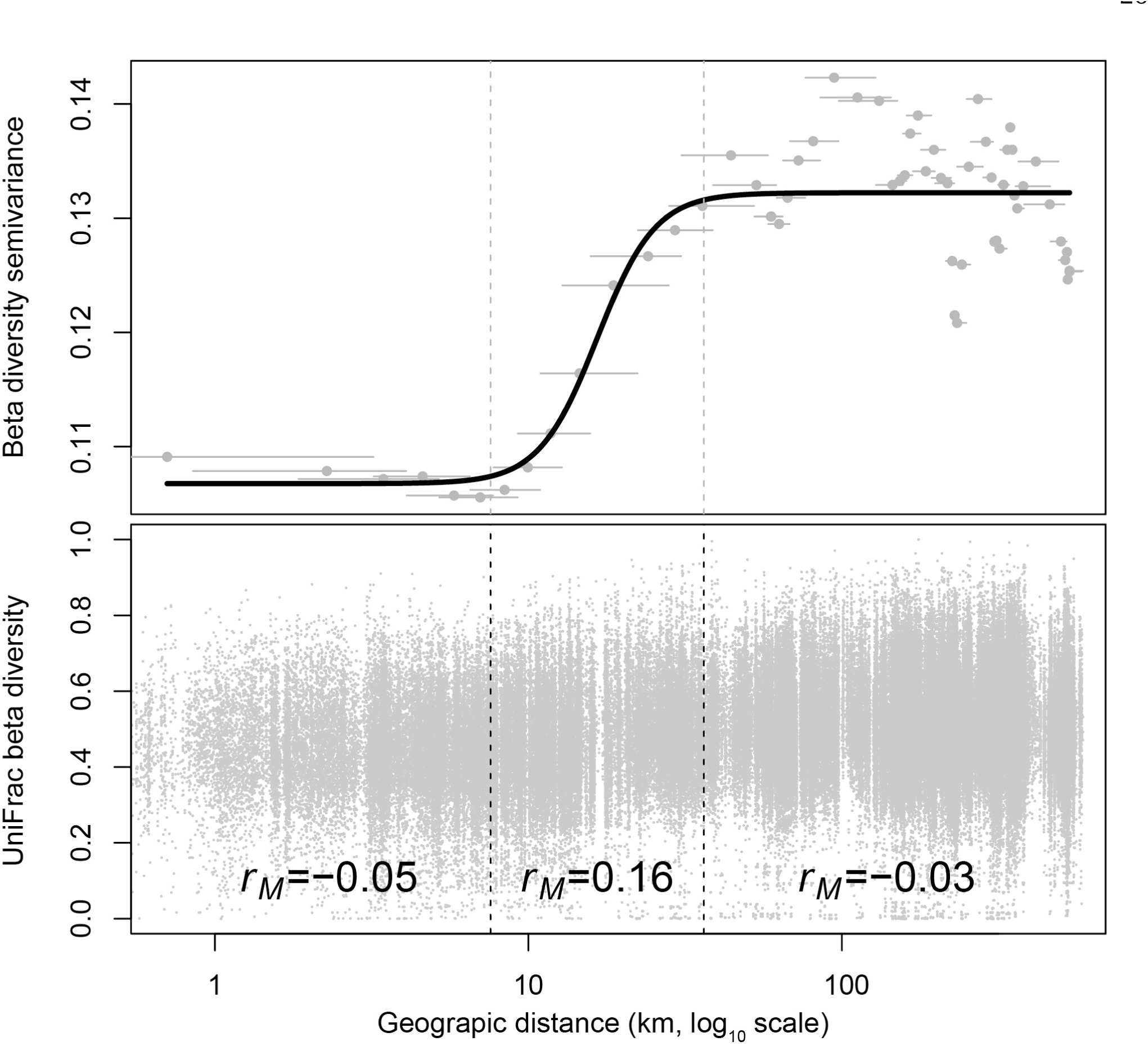
Semivariogram and Mantel correlations of FEF beta diversity. This semivariogram (Bachmaier & Backes, 2008) was constructed using geographic distance classes, each containing the same number of points and each sharing two-thirds of its points with the previous class. Dots indicate the mean geographic distance within each class, and horizontal lines indicate its geographic range. The black curve is a logistic function fit to the data. Increasing semivariance indicates spatial autocorrelation. The lag in spatial autocorrelation before 7.6 km (first dashed line) indicates a lack of spatial structuring, and that samples at that range may be drawn from the same community. The plateau and subsequent noise at 36.3 km (second dashed line) is called the “autocorrelation distance”, and is a typical feature of semivariogram analysis (King et al., 2010; Robeson et al., 2011). It indicates the distance after which spatial structuring becomes noisy or random, and geographic space is no longer a meaningful variable for community structure. The plot at bottom shows the data used to create the semivariogram. It also shows the Mantel correlation (*r*_*M*_) of the three distance classes, as determined by 99% intervals of the logistic fit’s derivative.

After GDM backward elimination, many variables were left as statistically significant: All *P*-values were the minimum possible *P*-value given that GDM’s backward elimination was run using 50 permutations. This means that GDM always predicted a smaller proportion of FEF beta diversity when any given predictor variable was permuted, except for sequencing run and transpiration which were eliminated this way. Host plant phylogenetic distance and evapotranspiration explained the most compositional dissimilarity in FEF communities, as given by their GDM coefficients (the maximum height of their splines) (Ferrier et al., 2007; Fitzpatrick et al., 2013) which were 0.090 and 0.078, respectively (Figure 4). These values indicate the relative importances of variables in the model. The shape of GDM *i*-splines reflects FEF community turnover over the range of differences in a predictor variable, and both host and evapotranspiration were related to FEF beta diversity over their entire ranges. Sample collection date was related to FEF beta diversity only at temporally proximate values, while solar radiation and NDVI were related to FEF beta diversity only at large differences (Figure 4). Although all variables shown in Figure 4 are statistically significant and the whole model fit nicely to the data with relatively symmetrical residuals (Figure 4, top), GDM was only able to predict 6% of FEF beta diversity. In contrast to the GDM fit using the UniFrac distance matrix, GDM run with Bray-Curtis dissimilarity fit poorly as a result of severe 1-inflation of distances (Supplementary figures S2 and S5).

**Figure 4:**
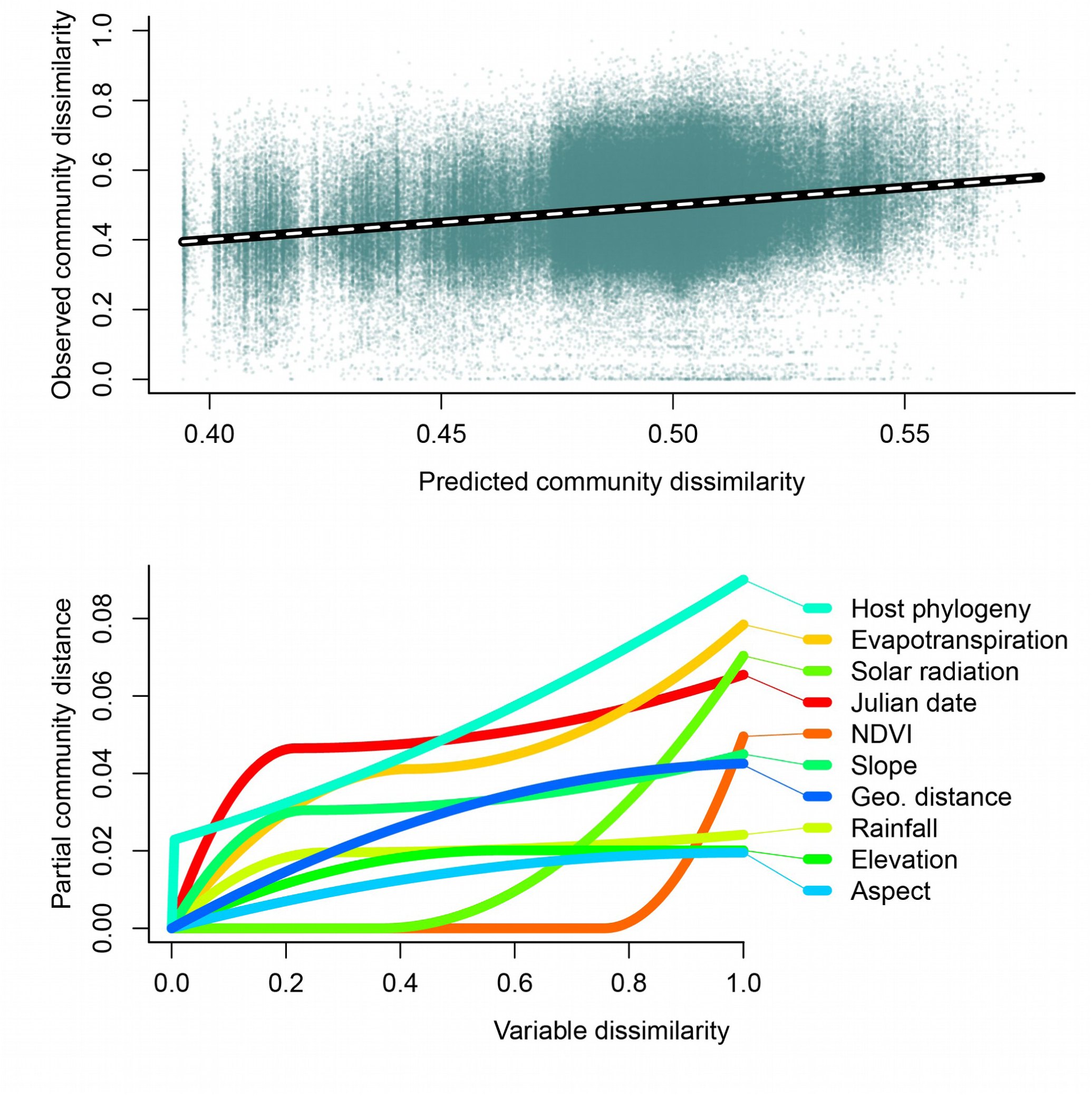
Model fit and coefficients for GDM model of FEF community dissimilarity. The observed community dissimilarity (UniFrac distance) between pairwise samples exhibited a linear but noisy relationship with the community dissimilarity predicted by the GDM model (top), which roughly corresponded to a 1:1 line (dashed line). In the bottom plot, GDM i-splines are shown for statistically significant predictor variables. Spline height indicates the relative importances of predictor variables, and the spline’s slope corresponds to the rate of change in compositional dissimilarity over the range of pairwise dissimilarities within the variable. Host phylogeny and Evapotranspiration were the strongest predictor variables in our analysis. These two variables were also the strongest correlates of beta diversity principal coordinate vectors in our confirmatory PCoA analysis.

Principal coordinate analysis (PCoA) was used to confirm GDM results using an alternative statistical approach. Similar to GDM, host (PCNM3 of host phylogenetic distance matrix) had the strongest relationship with FEF beta diverity with an *R*^2^ of 0.106 (*P* = 0.001), and evapotranspiration had the second strongest relationship with an R^2^ of 0.073 (*P* = 0.001). A table of PCoA *R*^2^ and *P* values is available as Supplementary Table S3.

### Bipartite network analysis

Each of the five islands we sampled showed a statistically significant (P < 0.001, Welch’s unequal variance *t*-test) pattern of FEF specialization (Figure 5), since FEF-island d’ values were higher than those generated using a null model (null d’ values centered around d’=0.25 for each island). Specialization in this case means that each island harbors more FEF OTUs that are uniquely associated with that island than would be expected by chance. We found that our sampling of host plant genera was significantly specialized to island as well (Supplementary figure S6), however plant-island d’ values were not significantly related to OTU-island d’ values (linear regression, *P* = 0.45). We also analyzed FEF-island specialization while keeping plant taxonomy identical across islands by randomly drawing a pre-set plant community from each island in each permutation of the d’ analysis. This analysis was done to remove the effect of plant community on FEF-island specialization. We still found significant specialization for all islands, although in this analysis all five islands had very similar d’ values (Supplementary figure S7).

**Figure 5:**
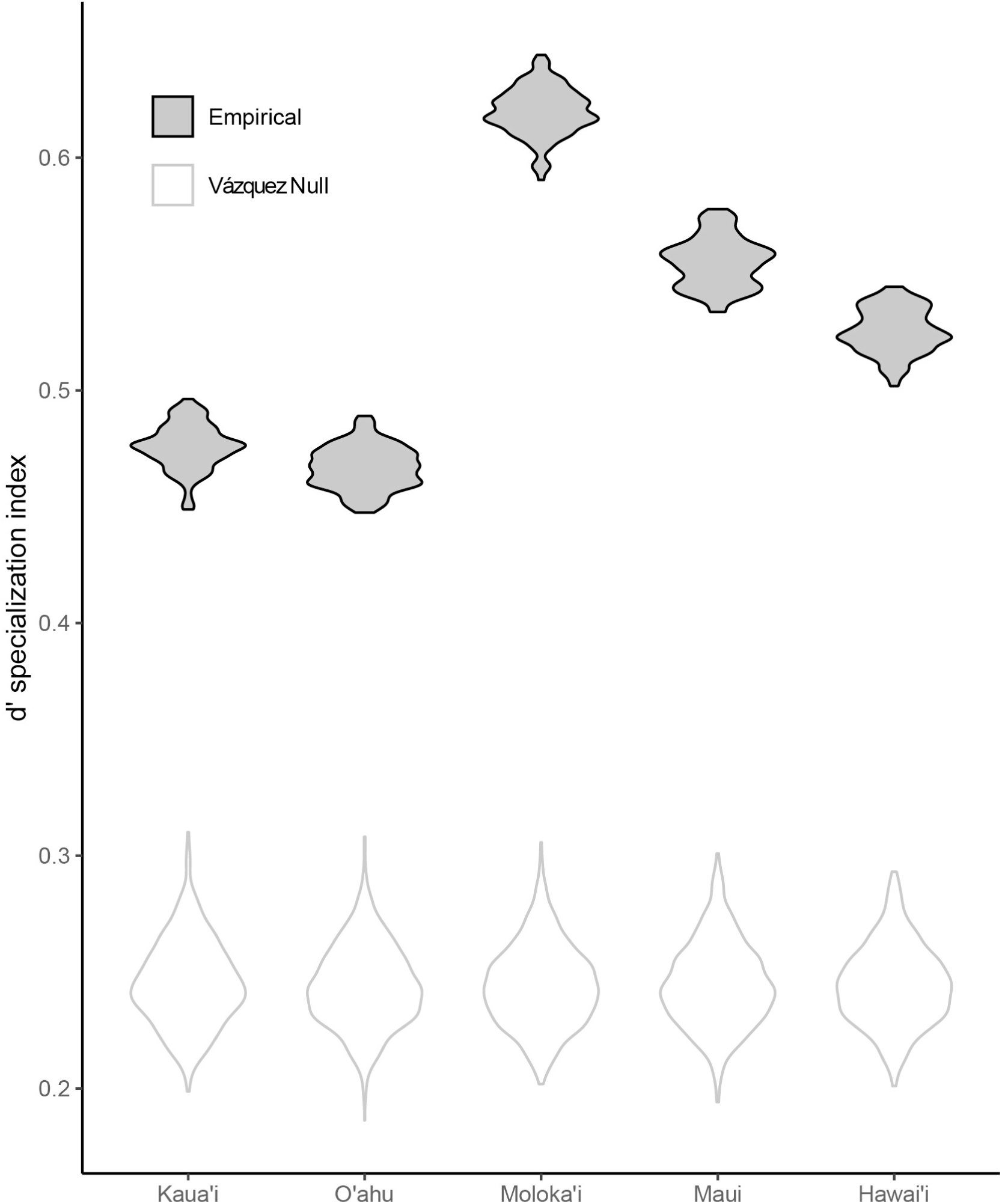
FEF specialization of each island. *d’* is a measure of how unique or cosmopolitan are the OTUs found on that island. In this violin plot, distributions shown in as filled violins with black borders are bootstrapped empirical *d’* values for each island. Unfilled distributions with a solid gray border are from the Vázquez null model, which is a fixed-connectance null model for bipartite networks. Welch’s unequal variance *t-*tests show that each island’s FEF community is significantly specialized as compared to the island null, meaning that FEF OTUs are more specific to their island of origin than expected by chance.

### Endophyte Taxonomic composition

At the Class level, FEF composition was consistent across islands, and dominated by Dothideomycetes and Sordariomycetes (members of the Pezizomycotina)(Supplementary figure S8). 12.14% of endophyte OTUs were unidentified at the class level (corresponding to 11.29% of rarefied sequence data), perhaps reflective of Hawaii’s isolation and comparative paucity of taxonomic research.

## DISCUSSION

The most striking pattern we found in our analysis of FEF communities in native Hawaiian plants was that evapotranspiration, a comparatively coarse measure of plant community traits at the landscape level, is meaningfully predictive for the community composition of microscopic fungi living within plant leaves. These results suggest that aside from environmental or host determinants, endophyte community composition results from a microbial “neighborhood”, in addition to the traits of individual hosts. Evapotranspiration was the second-most important variable in our analysis in terms of community composition (Figure 4); and it was more important in the model than elevation, which was measured at a much finer spatial resolution (Supplementary table S1). This was supported by our principal coordinate analysis as well. This result makes the interpretation of our hypothesis regarding the effect of climate on FEF community composition surprising, since previous studies (Coince et al., 2014; Zimmerman & Vitousek, 2012) suggested that temperature, elevation, and rainfall would be the most important factors structuring community composition instead, and evapotranspiration has not been previously considered as an important variable for FEF. While elevation (tightly correlated with temperature), host plant phylogeny, rainfall, and Julian date were significant explanatory variables in our GDM analysis, each of their effects were smaller than that of evapotranspiration (Figure 4).

Evapotranspiration encompasses both plant transpiration and the evaporation of water from soil and other surfaces, and both soil water content and stomatal conductance affect leaf interior moisture (Lambers, Chapin, & Pons, 2008). As such, evapotranspiration could drive FEF community composition by changing the leaf interior habitat, and thereby select for different FEF communities at high vs. low evapotranspiration. Indeed, evapotranspiration is strongly correlated with the moisture content of leaves (Lambers et al., 2008; Tardieu, Lafarge, & Simonneau, 1996). Alternatively, if plant stomata are open for longer periods, this may affect probabilities of different FEF successfully entering the leaf interior. To our knowledge, no studies have measured fungal community response to evapotranspiration, although associations between endophytic fungi and transpiration (a component of evapotranspiration) have been strong. In a grass system spanning 15 European countries, the response of endophytic fungi to a transpiration gradient was substantial (Lewis et al., 1997). Association between native FEF and transpiration has also been observed in *Theobroma cacao* (Arnold & Engelbrecht, 2007). In light of previous studies suggesting a link between fungal endophytes and evapotranspiration (Arnold & Engelbrecht, 2007; Kivlin et al., 2011; Lewis et al., 1997), our finding that evapotranspiration is a significant predictor of differences between FEF communities makes sense, and future physiological experiments at the scale of individual leaves might shine light on the nature of this interaction.

The association we found between FEF beta diversity and host plant phylogenetic distance was the most important predictor in our GDM model. Previous studies of FEF that have surveyed multiple host species, and have found that host identity is commonly associated with fungal endophyte community composition (Liu, Zhao, Wang, & Chen, 2019; U’Ren et al., 2012; Vincent et al., 2016), even among closely-related hosts (Christian et al., 2020), so our result was mostly expected. But unlike most previous studies that found host associations of FEF (although see (Davis & Shaw, 2008)), we used the phylogeny of host plants as an explanatory variable in place of their identity. Under the hypothesis that FEF communities are structured by host phylogenetic difference, more phylogenetically similar plants are expected to harbor similar FEF communities, and conversely, phylogenetically distant plants are expected to harbor less similar FEF communities. Following this hypothesis, in our analysis the GDM i-spline for host steadily increased across the whole range of differences in host phylogenetic distance (Figure 4), although the abrupt increase at low distances indicates rapid turnover in FEF community composition even with small changes in host phylogenetic distance. The significant association we observed from genus-level all the way up to higher taxonomic ranks agrees with a study of broad-scale host associations among tropical foliar fungal epiphytes, in which family level host taxonomy is most predictive (Kembel & Mueller, 2014).

This relationship between FEF beta diversity and host plant phylogeny could either mean that FEF, host plants, or both, exhibit a degree phylogenetic niche conservatism (Wiens et al., 2010). For example, FEF community preference may be phylogenetically conserved among closely related plant species, or perhaps host preference is conserved among closely related FEF. Patterns of phylosymbiosis are widespread throughout natural symbiotic systems, including plant-pathogen systems. However, phylogenetic signal can be lost in the presence of environmental factors that simultaneously impact symbiotic community composition (Brooks, Kohl, Brucker, van Opstal, & Bordenstein, 2016). Furthermore, in our analysis, host plant phylogeny may have been a more robust predictor of FEF community dissimilarity if our host plants had been classified to species level instead of genus level; however species level phylogenies are not fully resolved for many Hawaiian lineages, particularly those that have undergone rapid and recent adaptive radiations.

NDVI (normalized difference vegetation index) was a significant predictor of FEF community dissimilarity in our data set, suggesting that areas that are differentially vegetated harbor different communities of FEF. This difference may be related to the total percent land cover of vegetation, which is a component of NDVI (Purevdorj, Tateishi, Ishiyama, & Honda, 1998), or related to the type of plant cover, *i*.*e*. different plant community compositions (Lunetta, Knight, Ediriwickrema, Lyon, & Worthy, 2006), which is also encompassed within the NDVI metric. Indeed, foliar fungal communities potentially respond to both land cover and habitat type in the host plant *Quercus macrocarpa* (Jumpponen & Jones, 2009, 2010). This association between FEF and NDVI means that FEF conservation efforts would likely need to consider differential vegetation cover and urbanization at the landscape scale. The pixel size for the NDVI data we used was 250 m^2^ (Supplementary table S1), meaning that the value at any given location is an aggregate value for a large plant community. Thus, the proportion of FEF community dissimilarity that was significantly explained by NDVI in our analysis is related to the density and composition neighboring plant communities and not specifically the host individual, a dynamic suggested in others studies as well (Kato et al., 2017).

The other statistically significant drivers of FEF community composition (Figure 4) were mostly expected, particularly elevation (statistically significant in both GDM and PCoA analyses). This is similar to a result reported by Zimmerman and Vitousek (2012), who used PerMANOVA to test the effects of rainfall, elevation, and substrate age on FEF beta diversity patterns in *Metrosideros polymorpha* (‘Ō‘hia) trees. They found that elevation explained roughly 17% of compositional dissimilarity between FEF communities, varying slightly depending on which dissimilarity metric was used. That study also took place in Hawai‘i, and the area sampled overlaps partially with the area of Hawai‘i island that we sampled (Figure 1). Zimmerman and Vitousek (2012) also found a large effect of rainfall, which was also statistically significant in our GDM analysis. Rainfall is also a significant driver of FEF community structure in grasses (Giauque & Hawkes, 2013), although in a larger continental-scale analysis of cultured FEF isolates, rainfall was not a strong predictor of FEF diversity (U’Ren et al., 2012). Solar radiation was also a significant variable in our GDM and PCoA analyses, and was much more important in our GDM model than either elevation or rainfall. Indeed, previous research has shown that light levels are important for FEF colonization, likely because different intensities of light exposure can affect the chemical makeup of the phyllosphere (Bahnweg et al., 2005).

Geographic distance was of middling importance in our GDM analysis, especially since importance in GDM is calculated with all other variables accounted for. However, in our semivariogram and subsequent Mantel correlogram analyses, we found that FEF beta diversity is significantly spatially structured by geographic distance, especially between geographic distances of 7.6 and 36.3 km (Figure 3). Communities are not strongly spatially autocorrelated at distance of less than 7.6 km, and reach their autocorrelation range at 36.3 km. The lag before 7.6 km indicates that species are sampled from the same local community before that point (Robeson et al., 2011). After the curve plateaus around 36.3 km, FEF beta diversity reaches its “autocorrelation range”, after which autocorrelation becomes noisy. At distances greater than the autocorrelation range, communities are not spatially autocorrelated; *i*.*e*. community composition is no longer a function of geographic distance (King et al., 2010; Robeson et al., 2011). From our semivariogram we can qualitatively see that there may be spatial structuring at greater distances as well, perhaps up to 100 km, indicating that community structure is autocorrelated within islands, but not between them. Said another way, this indicates that the microbial “neighborhood” for FEF is at the sub-island scale. The Hawaiian islands’ volcanic origins might partially account for these patterns. Easterly trade winds and orographic lift drive much of the precipitation patterns in the archipelago, such that each island is neatly divided between comparatively hot and dry leeward, and cool wet windward sides. Nearly all island flora reflect this division, therefore it is unsurprising to find spatial autocorrelation at within, rather than among, island scales.

However, even though beta diversity may not be spatially autocorrelated at the inter-island scale, individual species may still be unique to islands. Said another way, islands may harbor unique cohorts of FEF in addition to cosmopolitan FEF, even though geographic space has little or no bearing on beta diversity at that scale. We tested this, and found a significant effect of FEF specialization to islands (Figure 5). Our bipartite network analysis showed that each island harbors a significant composition of fungi that are more regionally specific than would be expected due to chance alone. Beta diversity tools like GDM and our semivariogram are very different analytical methods than bipartite network analysis, and answer different questions. Our GDM analysis focused on patterns in FEF beta diversity, which is aggregate dissimilarity of community composition and does not take into account patterns at the OTU level, especially because of UniFrac’s down-weighting of phylogenetically similar OTUs. Our bipartite network analysis instead focused on the distribution of FEF OTUs within versus among islands, testing the extent to which a given island harbors OTUs that are exclusive to that island.

We presumed that the likeliest drivers of FEF-island specialization are differences in habitat, dispersal limitation among islands or differences in host availability (whether due to plant distributions or sampling bias). Since our beta diversity analysis revealed a significant effect of host plant phylogeny, because Hawaiian island floras are disharmonious (Ziegler, 2002), and because this disharmony was reflected in our sampling, differences in sampled plant hosts might account for specialization patterns. We used two separate analyses to evaluate whether this could account for specialization patterns alone. A bipartite analysis of plant-island specialization (Supplementary figure S6) indicated that in our experimental design, hosts are indeed specific to islands. However, we expected that if plant-island specialization bias within our experimental design was driving the FEF-island specialization pattern we observed, d’ values from the two would correlate strongly. There was no significant relationship between plant-island specialization and FEF-island specialization, and plant-island d’ values were much lower (less specialized) than those observed for OTUs. Further, when we restricted our analyses to only those plant genera that were sampled on all islands, the FEF specialization patterns remained.

How dispersal limitation shapes FEF-island specialization is an open question worthy of future research. There is some evidence for restricted gene flow among populations of a putative native mushroom, *Rhodocollybia laulaha* (Keirle, Avis, Feldheim, Hemmes, & Mueller, 2011) among islands, although these patterns might also be explained by selection. Similarly, the lag in our semivariogram (Figure 3) could be the distance at which dispersal limitation begins to structure FEF communities, or simply the distance at which un-measured environmental variables begin to vary enough to structure FEF communities. A long-term study of aerobiota at the summit of Hawaii’s peak, Mauna Loa, found evidence of likely long-distance dispersal via air mass (Tipton et al., 2019), although it is unclear how frequent and ecologically relevant such events are to endophyte systems. The epidemiology and rapid inter-island spread of fungal pathogens such as rapid ‘Ō‘hia death (*Ceratocystis* spp.) or ‘Ō‘hia rust (*Puccinia psidii*) suggest that at least some species are capable of crossing ocean channels, whether naturally or with the unwitting help of humans. Of course not all species are so easily dispersed, and the requisite and complex interaction between FEF, host and habitat likely reinforce island specialization patterns.

In summary, our analysis highlights that the various factors contributing to Hawaiian FEF community structure do so at the landscape scale. We analyzed these important plant symbionts across both a large geographic scale (Figure 1) and across a large host phylogenetic scale (80 plant genera), while using high-throughput sequencing to thoroughly inventory FEF community composition. We examined the effects of climate, geographic distance, and host identity using this system, and for the most part, found them to be strong predictors of differences in FEF communities between samples – even when the measurements were taken at relatively coarse resolution (Supplementary table S1). We found that elevation (Zimmerman & Vitousek, 2012), host plant phylogenetic difference (Cobian, Egan, & Amend, 2019; Huang et al., 2016; Kembel & Mueller, 2014; Massimo et al., 2015; Unterseher et al., 2012), geographic distance (U’Ren et al., 2012), and habitat type (Jumpponen & Jones, 2009, 2010; Kato et al., 2017) hypotheses were robust in our analysis at these large scales.

## AUTHOR CONTRIBUTIONS

JLD wrote the paper, performed bioinformatic and statistical analyses, and created figures. ASA and BAP conceived of the experiment. SOS, GC, and GLZ collected data and performed laboratory work. All authors contributed to the preparation of the manuscript.

## ACKNOWLEDGEMENTS

This work was supported by NSF award #1255972 to Amend, NSF award #1256128 to Perry and an NSF GRFP award to Cobian. The authors thank D Hemmes, J Adams, V Costello, E Datlof, S Walsh, K Kimball, M Purell, T Flynn, A Williams, R O’Rorke, S Perlman, P Biley and K Coelho for help in the field, and L Tooman for assistance in the lab. The authors additionally thank C Lozupone and J Foquier for helpful discussions. The authors declare that no conflict of interest exists.

## DATA ACCESSIBILITY STATEMENT

DNA sequence data have been submitted to the NCBI Sequence Read Archive (SRA) database under BioProject accession number PRJNA470970 (Darcy, Swift, et al., 2017). Computer code and input files that can be used to replicate this analysis are available at FigShare: https://figshare.com/s/34a59ebd8051589734c1 (temporary private link until publication). Additionally, a simple summary of the four most commonly sampled plant genera for each island is available as Supplementary table S4. DNA sequence data are available from both NCBI SRA and FigShare repositories.

## Notes

### Competing Interest Statement

The authors have declared no competing interest.

### Summary of Updates

changes for readability, methodological clarifications, minor edits.

