## Supplemental for "Fungal communities living within leaves of native Hawaiian dicots are structured by landscape-scale variables as well as by host plants"

#### Table of Contents:


| | |
|---|---|
| Figure S1 | Page 1 |
| Figure S2 | Page 2 |
| Figure S3 | Page 3 |
| Figure S4 | Page 4 |
| Figure S5 | Page 5 |
| Figure S6 | Page 6 |
| Figure S7 | Page 7 |
| Figure S8 | Page 8 |
| Table S1 | Page 9 |
| Table S2 | Page 10-23 |
| Table S3 | Page 24 |
| Table S4 | Page 25 |

#### Figure S1

Rarefaction curves. Top: accumulation of phylogenetic diversity (Faith, 1992) as rarefaction depth (D) is increased (sample is randomly downsampled to contain D sequences). In this figure, curves are binned by their phylogenetic diversity (Faith, 1992) at the rarefaction depth of D=1500 sequences per sample into 6 bins of even width (but varying numbers of curves within), to create averages (heavy lines). Binning was done because an average trend for samples with such high variance in diversity would not be meaningful. Bins were not used in the analysis of these data, and only serve to visually summarize rarefaction curves that overlap to such a high extent that without binning, individual curves for samples would be impossible to visually distinguish. Bottom: same as top plot but for number of OTUs observed instead of PD.

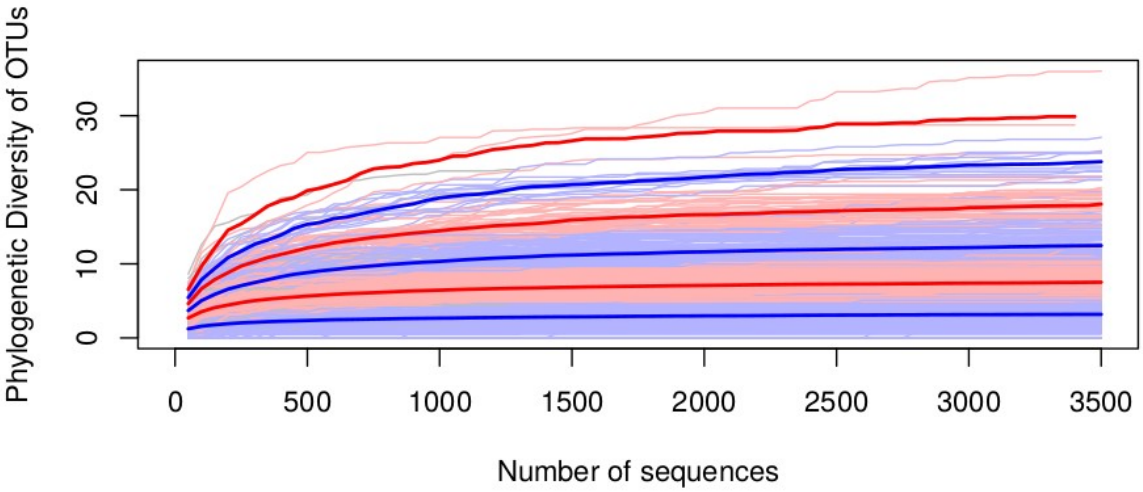


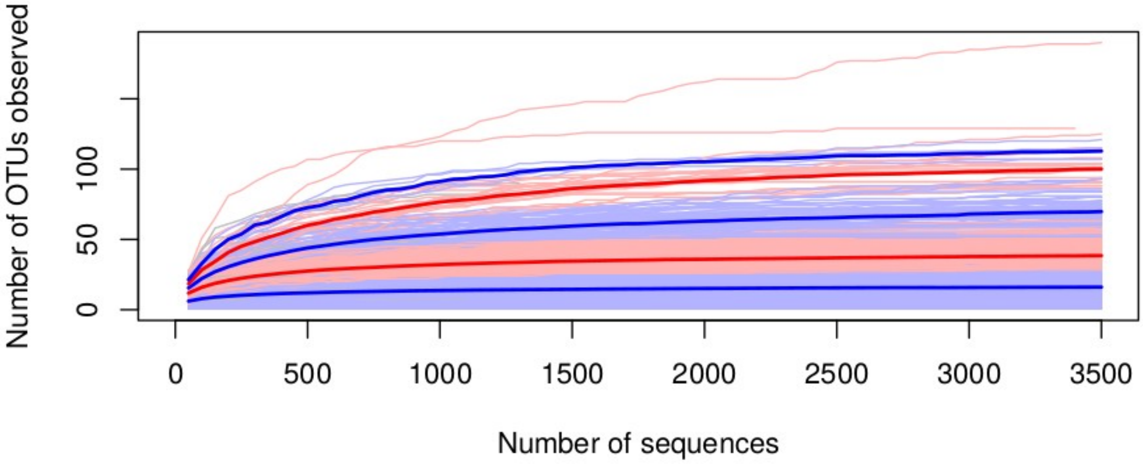

Figure S2

Relationship between UniFrac and Bray-Curtis dissimilarity matrices. Bray-Curtis dissimilarity (bottom) results in an inflation of 1-distances (max dissimilarity) because of sample pairs that share no OTUs (dashed lines corresponding to histograms represent the frequency of sample comparisons with a given dissimilarity bin). UniFrac (left) does not have this problem, because phylogenetic differences among OTUs are used instead of presence-absence differences (although both metrics are weighted by relative abundance).

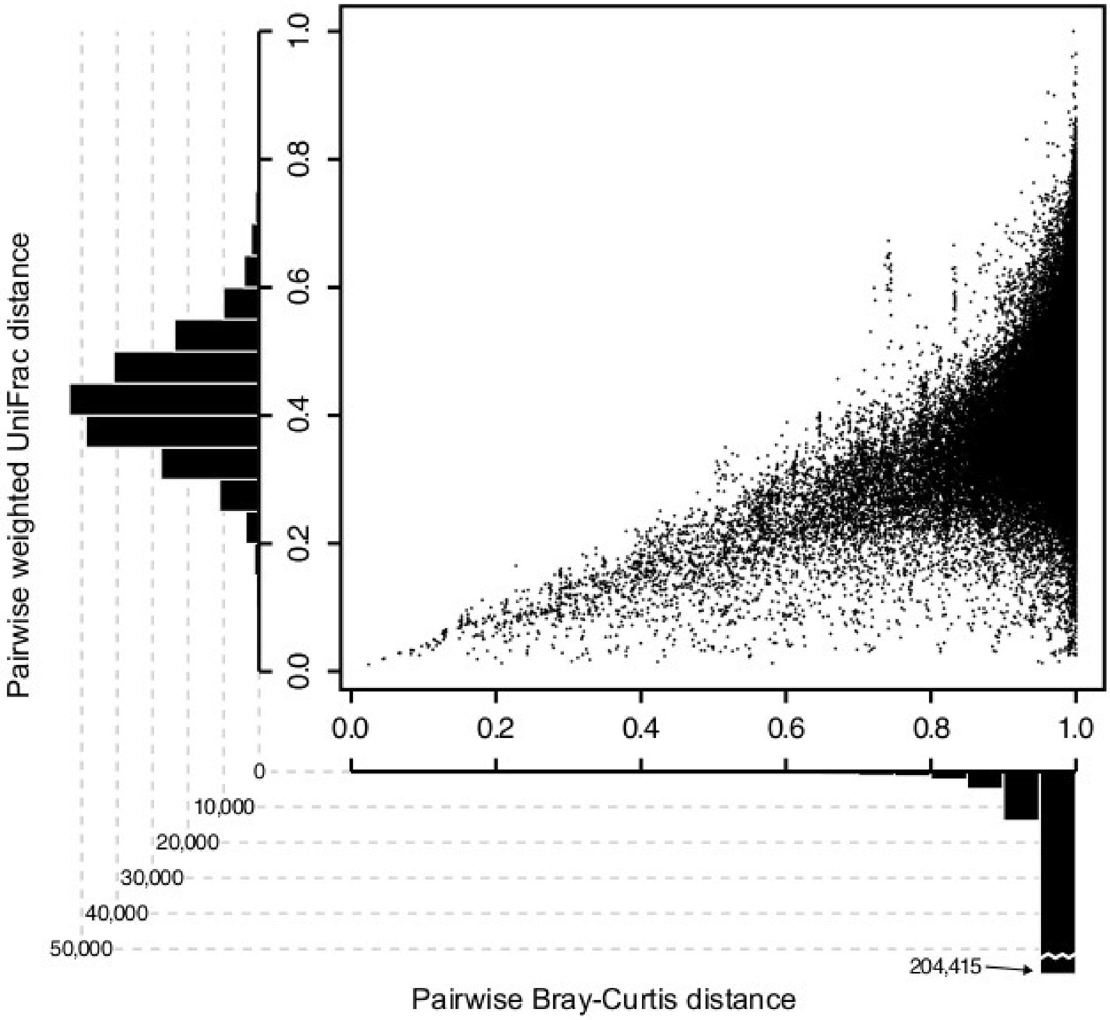

#### Figure S3

Predictor variable co-correlation. Some predictor variables in our analysis were strongly co-correlated, and as a result some were not included in the GDM model. In this figure, variable identity is shown in the diagonal, pairwise plots are shown in the lower triangle, and Spearman's rho correlation coefficient is shown in the upper triangle, in red if above 0.70.

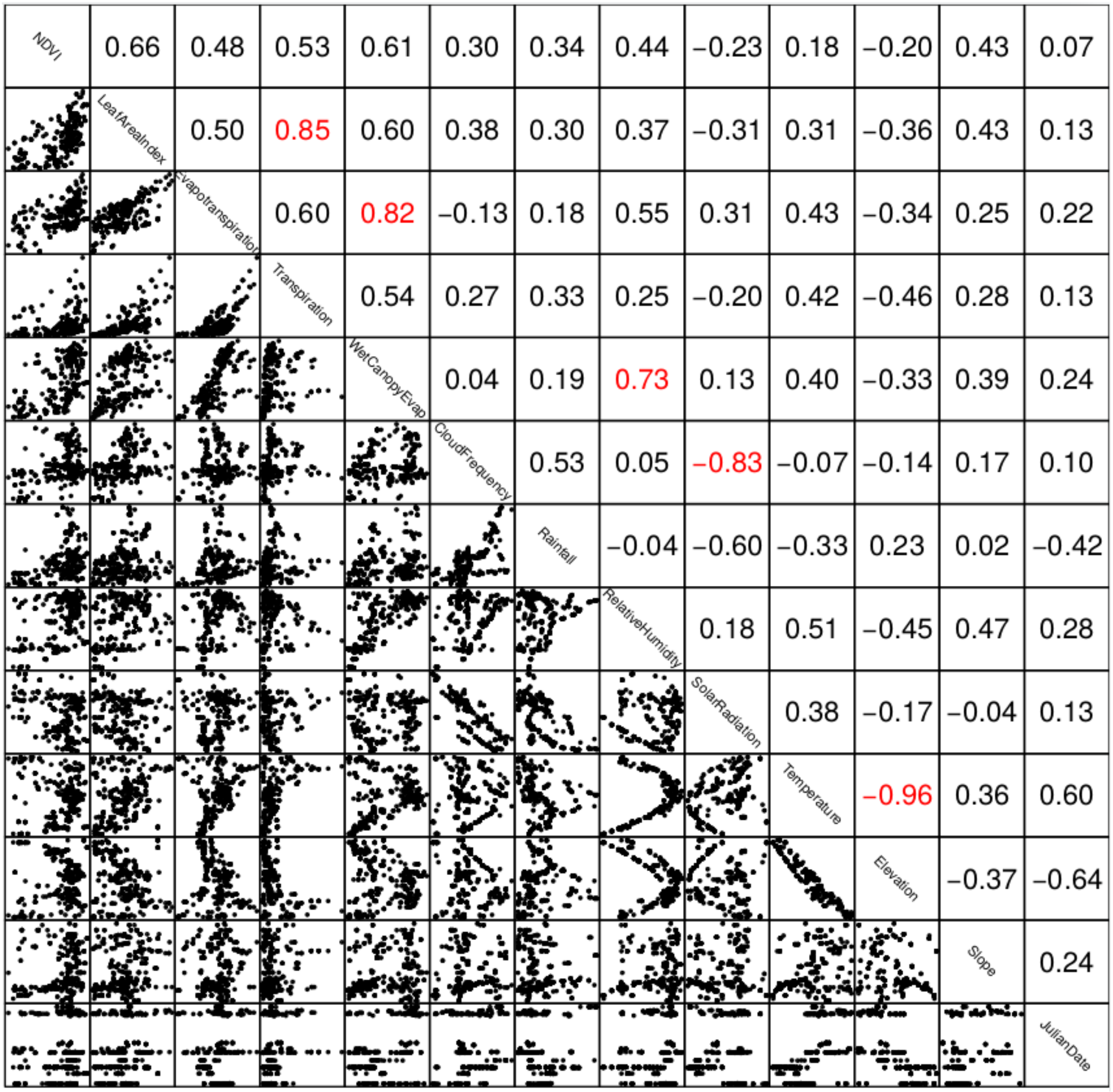

#### Figure S4

Host plant sampling by island. Black represents no sampling, yellow represents few individuals sampled, and red represents many individuals sampled. Phylogenetic relationships among host plants are shown at left.

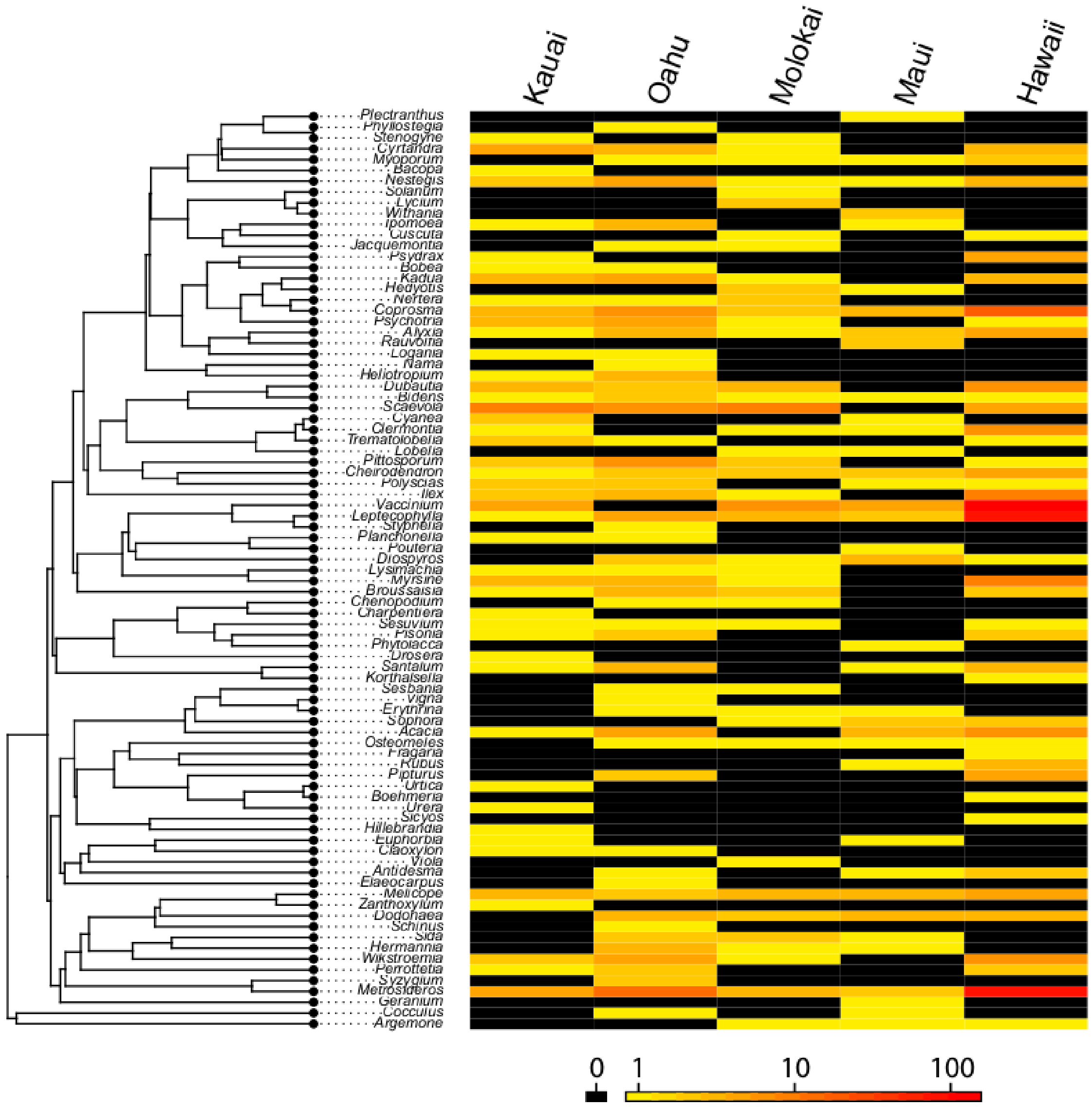

#### Figure S5

GDM model fitting comparison between different community distance matrices. Bray-Curtis produced a poor fit with heavily biased residuals, as a result of extreme 1-inflation (top panel). In comparison, the UniFrac matrix produced a model fit that is much more symmetrical about the 1:1 line (bottom, reproduced from Figure 3), similar to model fitting figures reported by other studies using GDM (e.g. Glassman et al. 2017, Fitzpatrick et al. 2013) and by the authors of the GDM R package in their vignette.

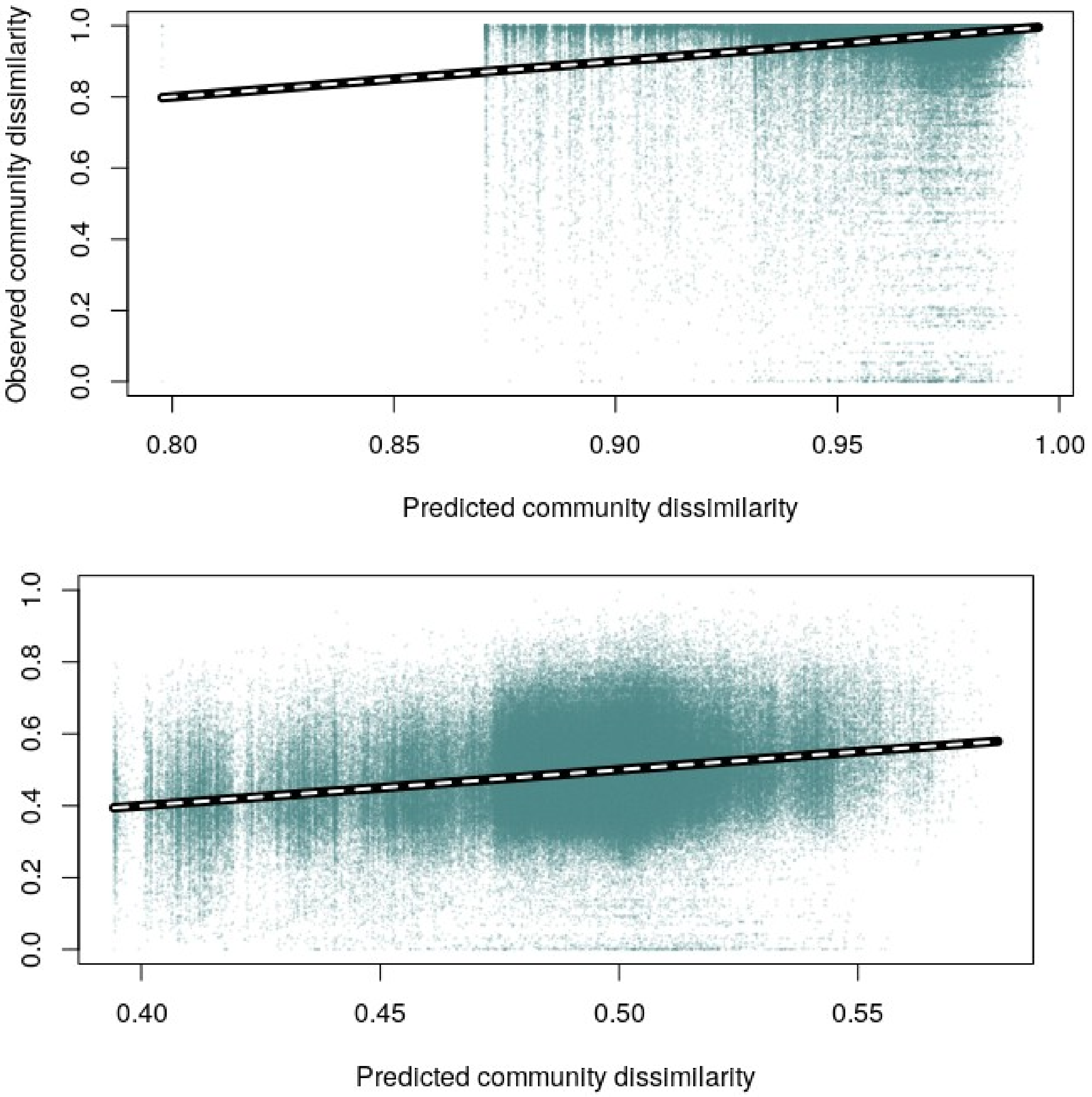

#### Figure S6

Specificity of plant genera to islands within our experiment. Empirical specificity values are shown as horizontal black lines, and null distributions are shown as gray violins. This analysis indicates that in our experimental design, hosts are indeed specific to islands. However, we expected that if host-island specificity bias within our experimental design was driving the OTU-island specificity pattern we observed, the two would strongly correlate. There was no significant relationship between host-island specificity and OTU-island specificity, and host-island *d'* values (shown here) were much lower than those observed for OTUs (Figure 4). So while part of the OTU specificity we observed may be due to bias in our sampling design, there is likely a significant pattern of OTU island specificity as well.

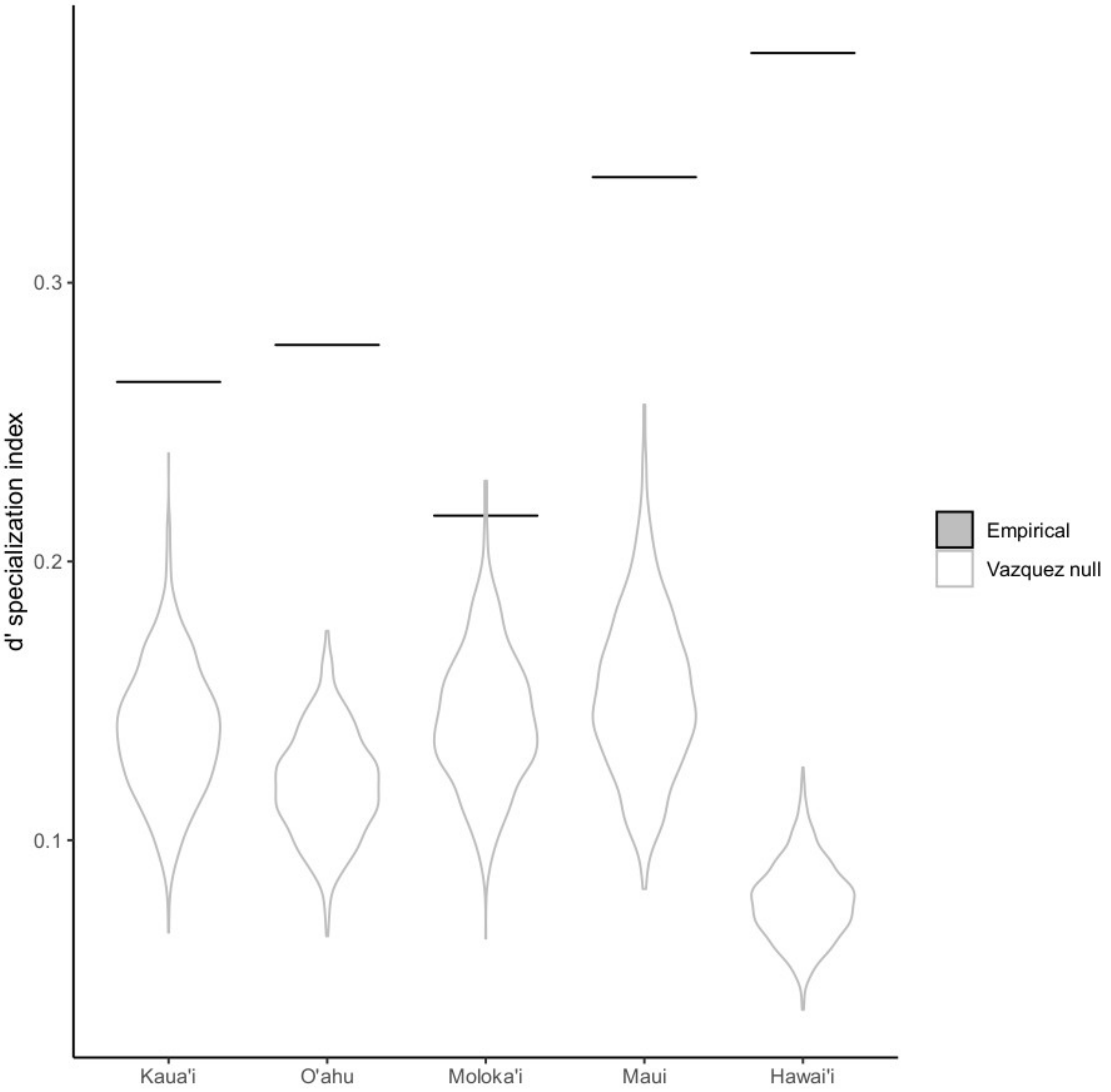

#### Figure S7

Island specificity with pre-set plant community. Because host plants were significantly specific to island within our experimental design (Fig. S4), we analyzed FEF island specificity using a subset of our data set where the plant community was fixed to be identical for each island. This fixed community was defined by the plant identification overlap between all five islands.

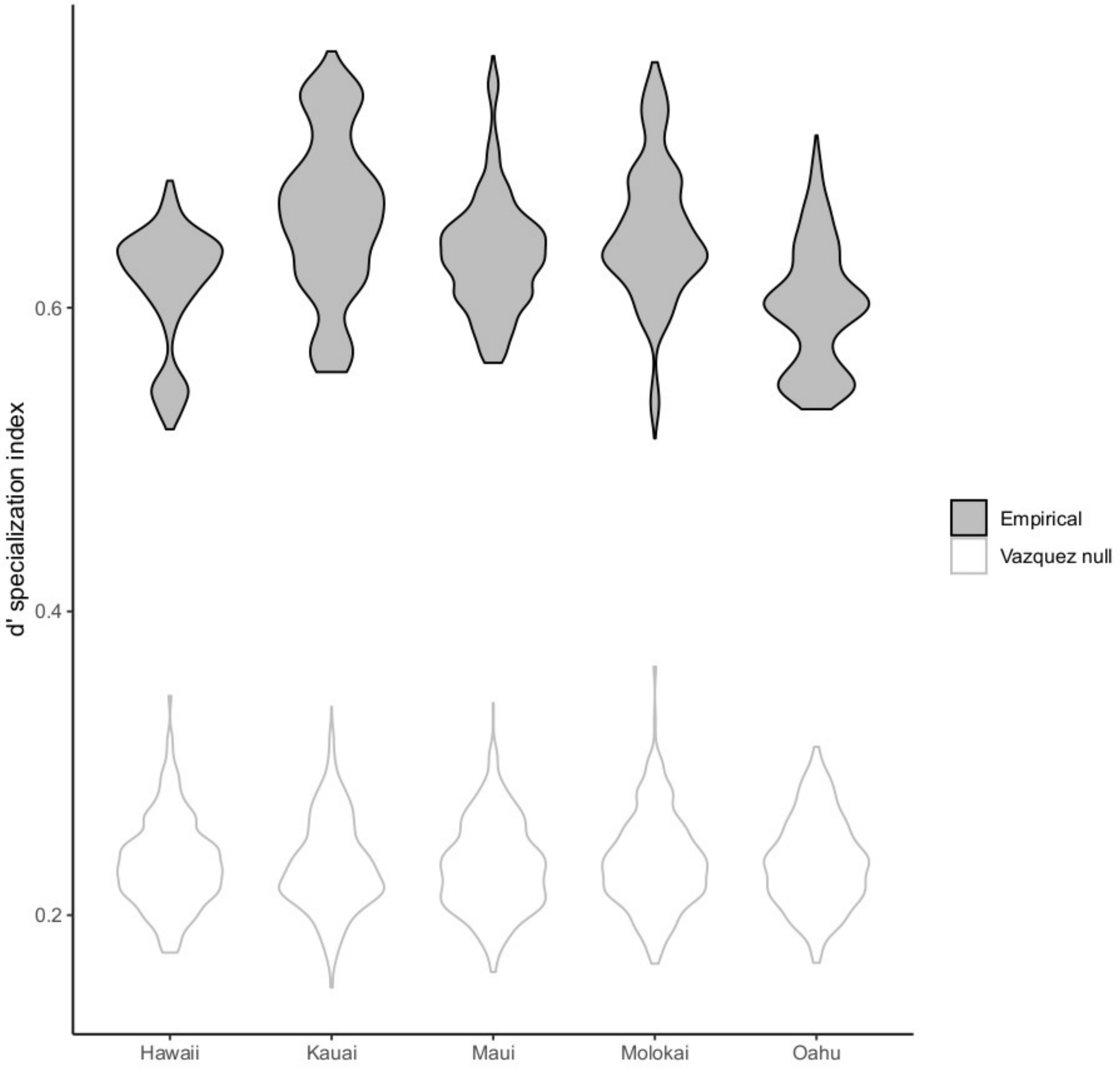

#### Figure S8

Class-level taxonomy of FEF by island. Subplots show the average proportion of reads per sample assigned to a given class. Error bars represent one SD. Classes with mean+SD < 0.02 are not shown. All islands were dominated by Dothideomycetes and Sordariomycetes (members of the Pezizomycotina). OTUs with unidentified class were still assigned to kingdom Fungi using the UNITE database as a reference.

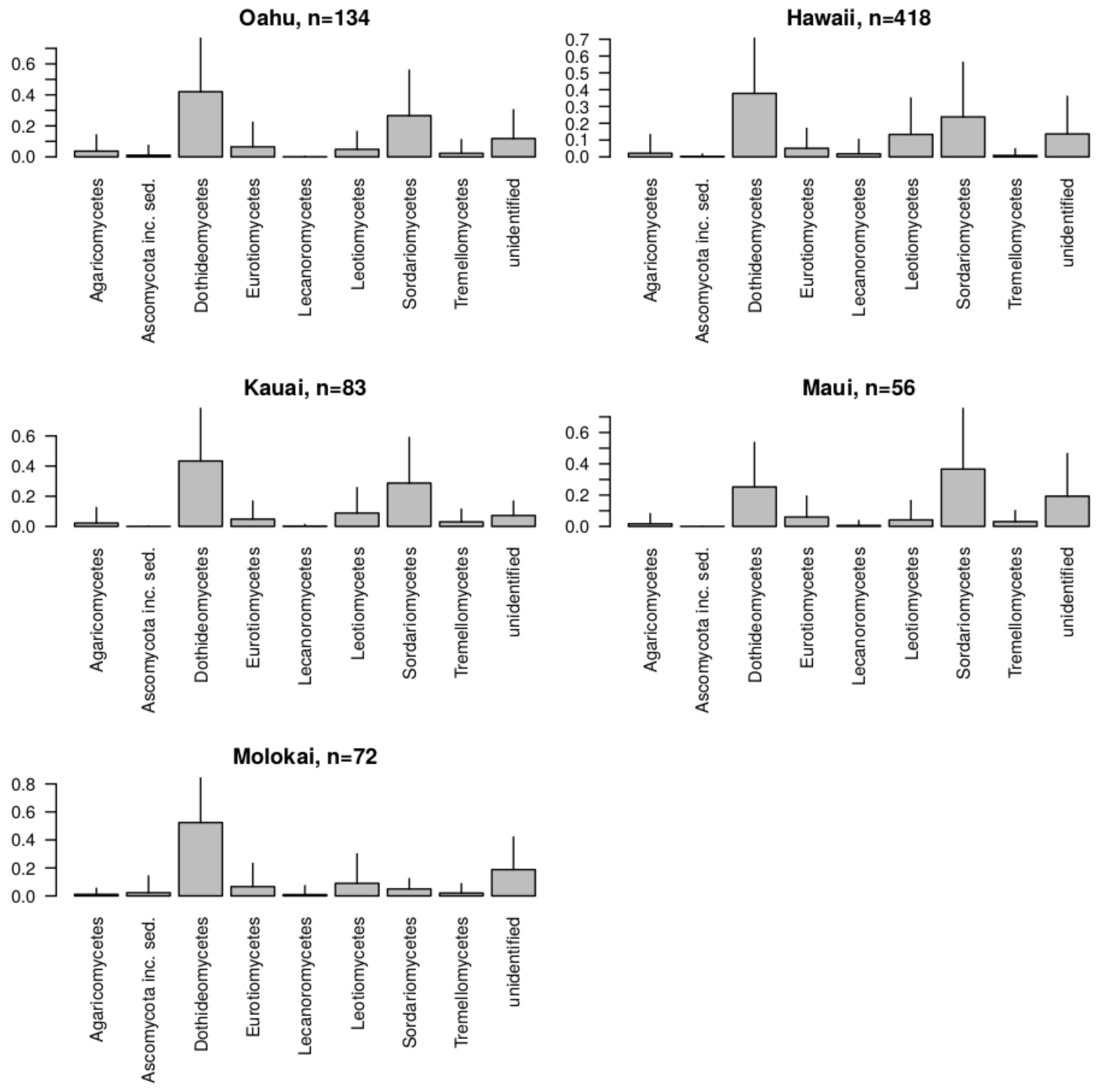

### Table S1

GIS data units, resolutions, and sources.

| Data type | Units | Pixel size | Source |
|---|---|---|---|
| Rainfall | mm*y-1 | 250 m2 | Rainfall atlas of Hawaiʻi |
| Evapotranspiration | mm*y-1 | 250 m2 | Evapotranspiration of Hawaiʻi |
| Leaf area index | index | 250 m2 | Evapotranspiration of Hawaiʻi |
| Wet Canopy Evaporation | mm*y-1 | 250 m2 | Evapotranspiration of Hawaiʻi |
| Transpiration | mm*y-1 | 250 m2 | Evapotranspiration of Hawaiʻi |
| Solar radiation | W/m2 | 250 m2 | Evapotranspiration of Hawaiʻi |
| Relative humidity | % | 250 m2 | Evapotranspiration of Hawaiʻi |
| Air temperature | °C | 250 m2 | Evapotranspiration of Hawaiʻi |
| Cloud frequency | ratio | 250 m2 | Evapotranspiration of Hawaiʻi |
| Elevation | m | 10.3 m2 | National elevation dataset |
| Slope | Degrees | 10.3 m2 | Calculated from elevation |
| Aspect | Degrees | 10.3 m2 | Calculated from elevation |
| NDVI | index | 250 m2 | Evapotranspiration of Hawaiʻi |

### Table S2

Geographic and taxonomic sample information. Plant species are putative because species level phylogenies are not fully resolved for many Hawaiian lineages, especially since many have undergone rapid and recent adaptive radiations. Many of the plants we collected lacked specific characters as well, making species-level identification uncertain.

| SampleID | Run | Latitude | Longitude | PlantGenus | PutativePlantSpecies | CollectionDate |
|---|---|---|---|---|---|---|
| BI036 | FEF2 | 19.406 | -155.253 | *Acacia* | *koa* | 07/21/14 |
| BI047 | FEF2 | 19.474 | -155.358 | *Acacia* | *koa* | 07/21/14 |
| BI069 | FEF2 | 19.687 | -155.466 | *Acacia* | *koa* | 07/23/14 |
| BI070 | FEF2 | 19.687 | -155.466 | *Acacia* | *koa* | 07/23/14 |
| BI126 | FEF2 | 19.664 | -155.388 | *Acacia* | *koa* | 07/25/14 |
| BI127 | FEF2 | 19.664 | -155.388 | *Acacia* | *koa* | 07/25/14 |
| BI143 | FEF2 | 19.649 | -155.372 | *Acacia* | *koa* | 07/25/14 |
| EKH5 | FEF2 | 21.439 | -158.095 | *Acacia* | *koa* | 09/22/15 |
| K0047 | FEF2 | 22.118 | -159.679 | *Acacia* | *koa* | 06/15/15 |
| UW03 | FEF2 | 20.773 | -156.235 | *Acacia* | *koa* | 11/06/14 |
| UW04 | FEF2 | 20.773 | -156.235 | *Acacia* | *koa* | 11/06/14 |
| wai19 | FEF2 | 20.799 | -156.253 | *Acacia* | *koa* | 11/04/14 |
| MC5 | FEF3 | 21.336 | -157.81 | *Acacia* | *koa* | 07/15/16 |
| MHL6 | FEF3 | 21.307 | -157.745 | *Acacia* | *koa* | 08/26/16 |
| PUPU1 | FEF3 | 21.641 | -158.018 | *Acacia* | *koa* | 08/03/16 |
| PUPU3 | FEF3 | 21.639 | -158.015 | *Acacia* | *koa* | 08/03/16 |
| BI055 | FEF2 | 20.113 | -155.759 | *Alyxia* | *oliviforme* | 07/22/14 |
| BI111 | FEF2 | 19.672 | -155.338 | *Alyxia* | *stellata* | 07/24/14 |
| BI154 | FEF2 | 20.09 | -155.737 | *Alyxia* | *stellata* | 07/26/14 |
| BI218 | FEF2 | 19.615 | -155.932 | *Alyxia* | *stellata* | 03/25/15 |
| EKH2 | FEF2 | 21.439 | -158.095 | *Alyxia* | *oliviforme* | 09/22/15 |
| EKH21 | FEF2 | 21.438 | -158.096 | *Alyxia* | *oliviforme* | 09/22/15 |
| K0058 | FEF2 | 22.15 | -159.623 | *Alyxia* | *oliviforme* | 06/16/15 |
| Kap12 | FEF2 | 20.933 | -156.633 | *Alyxia* | *stellata* | 06/11/15 |
| wai05 | FEF2 | 20.803 | -156.255 | *Alyxia* | *oliviforme* | 11/04/14 |
| KO17 | FEF3 | 21.352 | -157.793 | *Alyxia* | *stellata* | 08/11/16 |
| Mo074 | FEF3 | 21.107 | -156.901 | *Alyxia* | *stellata* | 07/19/16 |
| BI206 | FEF2 | 19.615 | -155.927 | *Antidesma* | *platyphyllum* | 03/25/15 |
| BI212 | FEF2 | 19.616 | -155.928 | *Antidesma* | *platyphyllum* | 03/25/15 |
| Kap19 | FEF2 | 20.933 | -156.63 | *Antidesma* | *pulvinatum* | 06/11/15 |
| MC9 | FEF3 | 21.335 | -157.81 | *Antidesma* | *platyphyllum* | 07/15/16 |
| BI068 | FEF2 | 19.687 | -155.466 | *Argemone* | *glauca* | 07/23/14 |
| Kan01 | FEF2 | 20.61 | -156.34 | *Argemone* | *glauca* | 06/10/15 |
| Mo044 | FEF3 | 21.091 | -156.928 | *Argemone* | *glauca* | 07/19/16 |
| K0100 | FEF2 | 22.223 | -159.416 | *Bacopa* | *monnieri* | 06/18/15 |
| BI219 | FEF2 | 19.615 | -155.932 | *Bidens* | *camphylotheca* | 03/25/15 |
| K0083 | FEF2 | 22.212 | -159.576 | *Bidens* | *forbsii* | 06/18/15 |
| Kap04 | FEF2 | 20.934 | -156.639 | *Bidens* | *micrantha* | 06/11/15 |
| L9 | FEF3 | 21.596 | -157.957 | *Bidens* | *macrocarpa* | 07/02/16 |
| Mo049 | FEF3 | 21.092 | -156.928 | *Bidens* | *wiebkei* | 07/19/16 |
| POAR6 | FEF3 | 21.527 | -157.92 | *Bidens* | *macrocarpa* | 10/02/16 |
| K0028 | FEF2 | 22.15 | -159.642 | *Bobea* | *brevipes* | 06/15/15 |
| POA20 | FEF3 | 21.534 | -157.924 | *Bobea* | *elatior* | 10/02/16 |
| BI229 | FEF2 | 19.616 | -155.931 | *Boehmeria* | *grandis* | 03/25/15 |
| BI064 | FEF2 | 20.114 | -155.76 | *Broussaisia* | *arguta* | 07/22/14 |
| BI162 | FEF2 | 20.091 | -155.738 | *Broussaisia* | *arguta* | 07/26/14 |
| K0022B | FEF2 | 22.151 | -159.643 | *Broussaisia* | *arguta* | 06/15/15 |
| SP18 | FEF2 | 21.512 | -158.137 | *Broussaisia* | *arguta* | 08/28/15 |

| | | | | | | |
|---|---|---|---|---|---|---|
| KO12 | FEF3 | 21.353 | -157.789 | *Broussaisia* | *arguta* | 08/11/16 |
| Mo023 | FEF3 | 21.118 | -156.905 | *Broussaisia* | *arguta* | 07/18/16 |
| Mo076 | FEF3 | 21.108 | -156.901 | *Broussaisia* | *arguta* | 07/19/16 |
| POAR4 | FEF3 | 21.526 | -157.918 | *Broussaisia* | *arguta* | 10/02/16 |
| K0084 | FEF2 | 22.212 | -159.576 | *Charpentiera* | *eliptica* | 06/18/15 |
| BI009 | FEF2 | 19.415 | -155.237 | *Cheirodendron* | *sp.* | 07/21/14 |
| BI056B | FEF2 | 20.113 | -155.759 | *Cheirodendron* | *trigynum* | 07/22/14 |
| BI110 | FEF2 | 19.672 | -155.338 | *Cheirodendron* | *sp.* | 07/24/14 |
| BI133 | FEF2 | 19.649 | -155.372 | *Cheirodendron* | *sp.* | 07/25/14 |
| BI147 | FEF2 | 20.09 | -155.737 | *Cheirodendron* | *sp.* | 07/26/14 |
| K0030 | FEF2 | 22.15 | -159.642 | *Cheirodendron* | *trigynum* | 06/15/15 |
| UW09 | FEF2 | 20.776 | -156.234 | *Cheirodendron* | *trigynum* | 11/06/14 |
| wai18 | FEF2 | 20.799 | -156.253 | *Cheirodendron* | *trigynum* | 11/04/14 |
| L3 | FEF3 | 21.596 | -157.958 | *Cheirodendron* | *trigynum* | 07/02/16 |
| Mo016 | FEF3 | 21.117 | -156.918 | *Cheirodendron* | *trigynum* | 07/18/16 |
| Mo073 | FEF3 | 21.105 | -156.902 | *Cheirodendron* | *trigynum* | 07/19/16 |
| POAR8 | FEF3 | 21.528 | -157.92 | *Cheirodendron* | *platyphyllum* | 10/02/16 |
| KEA13 | FEF3 | 21.572 | -158.21 | *Chenopodium* | *oahuense* | 08/18/16 |
| Mo047 | FEF3 | 21.092 | -156.928 | *Chenopodium* | *oahuense* | 07/19/16 |
| EKH29 | FEF2 | 21.439 | -158.094 | *Claoxylon* | *sandwicensis* | 09/22/15 |
| K0012 | FEF2 | 22.138 | -159.653 | *Claoxylon* | *sandwicensis* | 06/15/15 |
| BI005 | FEF2 | 19.414 | -155.238 | *Clermontia* | *sp.* | 07/21/14 |
| BI011 | FEF2 | 19.415 | -155.237 | *Clermontia* | *sp.* | 07/21/14 |
| BI062 | FEF2 | 20.113 | -155.76 | *Clermontia* | *parviflora* | 07/22/14 |
| BI167 | FEF2 | 20.091 | -155.74 | *Clermontia* | *sp.* | 07/26/14 |
| BI227 | FEF2 | 19.616 | -155.931 | *Clermontia* | *kohalae* | 03/25/15 |
| K0059 | FEF2 | 22.151 | -159.622 | *Clermontia* | *fauriei* | 06/16/15 |
| wai03 | FEF2 | 20.803 | -156.255 | *Clermontia* | *arborescens* | 11/04/14 |
| Mo043 | FEF3 | 21.126 | -156.919 | *Clermontia* | *sp.* | 07/18/16 |
| Kap06 | FEF2 | 20.934 | -156.638 | *Cocculus* | *orbiculatus* | 06/11/15 |
| LHL6 | FEF3 | 21.301 | -157.746 | *Cocculus* | *orbiculatus* | 08/21/16 |
| BI006 | FEF2 | 19.414 | -155.238 | *Coprosma* | *sp.* | 07/21/14 |
| BI056A | FEF2 | 20.113 | -155.759 | *Coprosma* | *sp.* | 07/22/14 |
| BI073 | FEF2 | 19.687 | -155.467 | *Coprosma* | *sp.* | 07/23/14 |
| BI079 | FEF2 | 19.687 | -155.469 | *Coprosma* | *sp.* | 07/23/14 |
| BI081 | FEF2 | 19.687 | -155.469 | *Coprosma* | *ernodeoides* | 07/23/14 |
| BI084 | FEF2 | 19.417 | -154.951 | *Coprosma* | *ochracea* | 07/24/14 |
| BI104 | FEF2 | 19.672 | -155.338 | *Coprosma* | *sp.* | 07/24/14 |
| BI105 | FEF2 | 19.672 | -155.338 | *Coprosma* | *ernodeoides* | 07/24/14 |
| BI114 | FEF2 | 19.672 | -155.338 | *Coprosma* | *ochracea* | 07/24/14 |
| BI120 | FEF2 | 19.676 | -155.384 | *Coprosma* | *ernodeoides* | 07/25/14 |
| BI124 | FEF2 | 19.673 | -155.385 | *Coprosma* | *sp.* | 07/25/14 |
| BI151 | FEF2 | 20.09 | -155.737 | *Coprosma* | *sp.* | 07/26/14 |
| BI166 | FEF2 | 20.091 | -155.74 | *Coprosma* | *longerleaf* | 07/26/14 |
| BI205 | FEF2 | 19.615 | -155.927 | *Coprosma* | *arychnospra* | 03/25/15 |
| BI208 | FEF2 | 19.615 | -155.927 | *Coprosma* | *menzisii* | 03/25/15 |
| BI221 | FEF2 | 19.615 | -155.932 | *Coprosma* | *sp.* | 03/25/15 |
| EKH16 | FEF2 | 21.439 | -158.097 | *Coprosma* | *sp.* | 09/22/15 |
| K0015 | FEF2 | 22.138 | -159.653 | *Coprosma* | *kauensis* | 06/15/15 |
| K0025 | FEF2 | 22.151 | -159.643 | *Coprosma* | *waimaeae* | 06/15/15 |
| K0066 | FEF2 | 22.152 | -159.616 | *Coprosma* | *eliptica* | 06/16/15 |
| SP03 | FEF2 | 21.512 | -158.136 | *Coprosma* | *longiflora* | 08/28/15 |
| SP04 | FEF2 | 21.512 | -158.136 | *Coprosma* | *longiflora* | 08/28/15 |
| SP07 | FEF2 | 21.512 | -158.136 | *Coprosma* | *longiflora* | 08/28/15 |
| SP08 | FEF2 | 21.512 | -158.136 | *Coprosma* | *longiflora* | 08/28/15 |
| SP13 | FEF2 | 21.512 | -158.136 | *Coprosma* | *longiflora* | 08/28/15 |
| SP17 | FEF2 | 21.512 | -158.136 | *Coprosma* | *longiflora* | 08/28/15 |
| UW06 | FEF2 | 20.773 | -156.235 | *Coprosma* | *arenioides* | 11/06/14 |
| UW16 | FEF2 | 20.776 | -156.234 | *Coprosma* | *montana* | 11/06/14 |

| | | | | | | |
|---|---|---|---|---|---|---|
| wai04 | FEF2 | 20.803 | -156.255 | *Coprosma* | *foliosa* | 11/04/14 |
| Mo009 | FEF3 | 21.119 | -156.926 | *Coprosma* | *sp.* | 07/18/16 |
| Mo012 | FEF3 | 21.117 | -156.919 | *Coprosma* | *sp.* | 07/18/16 |
| BI090 | FEF2 | 19.448 | -154.862 | *Cuscuta* | *sandwichiana* | 07/24/14 |
| Mo092 | FEF3 | 21.2 | -157.158 | *Cuscuta* | *sandwichiana* | 07/20/16 |
| K0062 | FEF2 | 22.154 | -159.618 | *Cyanea* | *hirtella* | 06/16/15 |
| K0094 | FEF2 | 22.216 | -159.579 | *Cyanea* | *coriaceae* | 06/18/15 |
| UW12 | FEF2 | 20.776 | -156.234 | *Cyanea* | *linearis* | 11/06/14 |
| BI015 | FEF2 | 19.415 | -155.237 | *Cyrtandra* | *sp.* | 07/21/14 |
| BI087 | FEF2 | 19.417 | -154.951 | *Cyrtandra* | *platyphylla* | 07/24/14 |
| BI164 | FEF2 | 20.091 | -155.739 | *Cyrtandra* | *sp.* | 07/26/14 |
| EKH14 | FEF2 | 21.439 | -158.096 | *Cyrtandra* | *waianaensis* | 09/22/15 |
| K0035A | FEF2 | 22.148 | -159.636 | *Cyrtandra* | *platyphylla* | 06/15/15 |
| K0056 | FEF2 | 22.15 | -159.623 | *Cyrtandra* | *longifolia* | 06/16/15 |
| K0089 | FEF2 | 22.209 | -159.579 | *Cyrtandra* | *wainihaensis* | 06/18/15 |
| K0091A | FEF2 | 22.209 | -159.579 | *Cyrtandra* | *confertiflora* | 06/18/15 |
| K0091B | FEF2 | 22.209 | -159.579 | *Cyrtandra* | *confertiflora* | 06/18/15 |
| KO13 | FEF3 | 21.353 | -157.789 | *Cyrtandra* | *paludosa* | 08/11/16 |
| MC8 | FEF3 | 21.335 | -157.811 | *Cyrtandra* | *sp.* | 07/15/16 |
| Mo026 | FEF3 | 21.118 | -156.902 | *Cyrtandra* | *sp.* | 07/18/16 |
| BI239 | FEF2 | 19.112 | -155.823 | *Diospyros* | *sandwicensis* | 03/25/15 |
| Kan05 | FEF2 | 20.61 | -156.34 | *Diospyros* | *sandwicensis* | 06/10/15 |
| Kan18 | FEF2 | 20.62 | -156.35 | *Diospyros* | *sandwicensis* | 06/10/15 |
| Kap10 | FEF2 | 20.934 | -156.637 | *Diospyros* | *sandwicensis* | 06/11/15 |
| HL2 | FEF3 | 21.302 | -157.745 | *Diospyros* | *sandwicensis* | 07/21/16 |
| Mo052 | FEF3 | 21.1 | -156.914 | *Diospyros* | *sandwicensis* | 07/19/16 |
| BI034 | FEF2 | 19.409 | -155.254 | *Dodonaea* | *viscosa* | 07/21/14 |
| BI080 | FEF2 | 19.687 | -155.469 | *Dodonaea* | *viscosa* | 07/23/14 |
| BI122 | FEF2 | 19.675 | -155.384 | *Dodonaea* | *viscosa* | 07/25/14 |
| Kan13 | FEF2 | 20.61 | -156.35 | *Dodonaea* | *viscosa* | 06/10/15 |
| Kap05 | FEF2 | 20.934 | -156.638 | *Dodonaea* | *viscosa* | 06/11/15 |
| UW18 | FEF2 | 20.766 | -156.257 | *Dodonaea* | *viscosa* | 11/06/14 |
| KEA4 | FEF3 | 21.563 | -158.211 | *Dodonaea* | *viscosa* | 08/18/16 |
| KEA7 | FEF3 | 21.569 | -158.21 | *Dodonaea* | *viscosa* | 08/18/16 |
| LHL2 | FEF3 | 21.301 | -157.745 | *Dodonaea* | *viscosa* | 08/21/16 |
| Mo007 | FEF3 | 21.119 | -156.93 | *Dodonaea* | *viscosa* | 07/18/16 |
| Mo048 | FEF3 | 21.092 | -156.928 | *Dodonaea* | *viscosa* | 07/19/16 |
| K0109 | FEF2 | 21.975 | -159.507 | *Drosera* | *anglica* | 06/19/15 |
| BI028 | FEF2 | 19.411 | -155.251 | *Dubautia* | *sp.* | 07/21/14 |
| BI030 | FEF2 | 19.411 | -155.251 | *Dubautia* | *sp.* | 07/21/14 |
| BI043 | FEF2 | 19.42 | -155.29 | *Dubautia* | *ciliolata* | 07/21/14 |
| BI093 | FEF2 | 19.676 | -155.329 | *Dubautia* | *ciliolata* | 07/24/14 |
| BI117 | FEF2 | 19.672 | -155.336 | *Dubautia* | *ciliolata* | 07/24/14 |
| BI130 | FEF2 | 19.66 | -155.386 | *Dubautia* | *ciliolata* | 07/25/14 |
| K0039 | FEF2 | 22.149 | -159.636 | *Dubautia* | *laevigata* | 06/15/15 |
| K0054 | FEF2 | 22.15 | -159.623 | *Dubautia* | *raillardiodes* | 06/16/15 |
| K0119 | FEF2 | 21.984 | -159.501 | *Dubautia* | *imbricata* | 06/19/15 |
| SP09 | FEF2 | 21.512 | -158.136 | *Dubautia* | *laxa* | 08/28/15 |
| KO2 | FEF3 | 21.355 | -157.788 | *Dubautia* | *laxa* | 08/11/16 |
| Mo034 | FEF3 | 21.119 | -156.9 | *Dubautia* | *laxa* | 07/18/16 |
| Mo069 | FEF3 | 21.102 | -156.906 | *Dubautia* | *linearis* | 07/19/16 |
| Mo071 | FEF3 | 21.105 | -156.902 | *Dubautia* | *plantaginea* | 07/19/16 |
| POAR12 | FEF3 | 21.532 | -157.921 | *Elaeocarpus* | *bifidus* | 10/02/16 |
| Kan15 | FEF2 | 20.61 | -156.35 | *Erythrina* | *sandwicensis* | 06/10/15 |
| KEA11 | FEF3 | 21.572 | -158.211 | *Erythrina* | *sandwicensis* | 08/18/16 |
| Mo081 | FEF3 | 21.083 | -156.935 | *Erythrina* | *sandwicensis* | 07/19/16 |
| K0105 | FEF2 | 21.975 | -159.508 | *Euphorbia* | *sprsiflora* | 06/19/15 |
| Kap11 | FEF2 | 20.934 | -156.637 | *Euphorbia* | *anotiana* | 06/11/15 |
| BI027 | FEF2 | 19.412 | -155.251 | *Fragaria* | *chiloensis* | 07/21/14 |

| | | | | | | |
|---|---|---|---|---|---|---|
| UW17 | FEF2 | 20.768 | -156.238 | *Geranium* | *cuneatum* | 11/06/14 |
| wai17 | FEF2 | 20.799 | -156.253 | *Hedyotis* | *hillebrandii* | 11/04/14 |
| Mo011 | FEF3 | 21.117 | -156.919 | *Hedyotis* | *sp.* | 07/18/16 |
| Mo058 | FEF3 | 21.1 | -156.914 | *Hedyotis* | *hillebrandii* | 07/19/16 |
| K0077 | FEF2 | 21.893 | -159.405 | *Heliotropium* | *anomalum* | 06/17/15 |
| Maka03 | FEF2 | 21.289 | -157.666 | *Heliotropium* | *curassavicum* | 07/21/15 |
| Maka11 | FEF2 | 21.293 | -157.659 | *Heliotropium* | *anomalum* | 07/21/15 |
| Maka15 | FEF2 | 21.316 | -157.663 | *Heliotropium* | *anomalum* | 07/21/15 |
| Kan02 | FEF2 | 20.61 | -156.34 | *Waltheria* | *indica* | 06/10/15 |
| Maka08P4 | FEF2 | 21.292 | -157.662 | *Waltheria* | *indica* | 07/21/15 |
| KEA3 | FEF3 | 21.563 | -158.211 | *Waltheria* | *indica* | 08/18/16 |
| Mo051 | FEF3 | 21.1 | -156.914 | *Waltheria* | *indica* | 07/19/16 |
| WS1 | FEF3 | 21.352 | -158.13 | *Waltheria* | *indica* | 07/30/16 |
| K0008 | FEF2 | 22.131 | -159.655 | *Hillebrandia* | *sandwicensis* | 06/15/15 |
| X69cv07 | FEF2 | 21.515 | -158.161 | *Ilex* | *anomala* | 09/08/15 |
| BI002 | FEF2 | 19.414 | -155.239 | *Ilex* | *ambigua* | 07/21/14 |
| BI026 | FEF2 | 19.412 | -155.251 | *Ilex* | *ambigua* | 07/21/14 |
| BI054 | FEF2 | 20.113 | -155.759 | *Ilex* | *anomala* | 07/22/14 |
| BI100 | FEF2 | 19.674 | -155.33 | *Ilex* | *anomala* | 07/24/14 |
| BI115 | FEF2 | 19.672 | -155.338 | *Ilex* | *anomala* | 07/24/14 |
| BI132 | FEF2 | 19.652 | -155.376 | *Ilex* | *anomala* | 07/25/14 |
| BI160 | FEF2 | 20.091 | -155.738 | *Ilex* | *anomala* | 07/26/14 |
| BI230 | FEF2 | 19.615 | -155.931 | *Ilex* | *anomola* | 03/25/15 |
| K0027A | FEF2 | 22.15 | -159.642 | *Ilex* | *anomala* | 06/15/15 |
| K0027B | FEF2 | 22.15 | -159.642 | *Ilex* | *anomala* | 06/15/15 |
| SP19 | FEF2 | 21.512 | -158.137 | *Ilex* | *anomala* | 08/28/15 |
| L6 | FEF3 | 21.596 | -157.958 | *Ilex* | *anomala* | 07/02/16 |
| Mo018 | FEF3 | 21.118 | -156.905 | *Ilex* | *anomala* | 07/18/16 |
| K0072 | FEF2 | 21.892 | -159.411 | *Ipomoea* | *pescaprae* | 06/17/15 |
| Kan11 | FEF2 | 20.61 | -156.34 | *Ipomoea* | *tuboides* | 06/10/15 |
| Maka06 | FEF2 | 21.292 | -157.662 | *Ipomoea* | *sp.* | 07/21/15 |
| Maka17 | FEF2 | 21.316 | -157.663 | *Ipomoea* | *sp.* | 07/21/15 |
| NS5 | FEF3 | 21.581 | -158.207 | *Ipomoea* | *pes-caprae* | 08/03/16 |
| Maka18 | FEF2 | 21.315 | -157.662 | *Jacquemontia* | *sandwicensis* | 07/21/15 |
| Mo082 | FEF3 | 21.2 | -157.157 | *Jacquemontia* | *sandwicensis* | 07/20/16 |
| X69cv13 | FEF2 | 21.515 | -158.161 | *Kadua* | *terminalis* | 09/08/15 |
| BI049 | FEF2 | 20.113 | -155.758 | *Kadua* | *sp.* | 07/22/14 |
| BI215 | FEF2 | 19.617 | -155.929 | *Kadua* | *kiminalis* | 03/25/15 |
| BI224 | FEF2 | 19.615 | -155.932 | *Kadua* | *kiminalis* | 03/25/15 |
| EKH22 | FEF2 | 21.438 | -158.096 | *Kadua* | *terminalis* | 09/22/15 |
| K0024 | FEF2 | 22.151 | -159.643 | *Kadua* | *affinis* | 06/15/15 |
| K0026 | FEF2 | 22.15 | -159.642 | *Kadua* | *foggiana* | 06/15/15 |
| K0088 | FEF2 | 22.21 | -159.578 | *Kadua* | *accuminata* | 06/18/15 |
| HL4 | FEF3 | 21.317 | -157.743 | *Kadua* | *affinis* | 07/21/16 |
| KO11 | FEF3 | 21.354 | -157.788 | *Kadua* | *fosbergii* | 08/11/16 |
| Mo075 | FEF3 | 21.108 | -156.901 | *Kadua* | *sp.* | 07/19/16 |
| POAR15 | FEF3 | 21.533 | -157.921 | *Kadua* | *sp.* | 10/02/16 |
| BI155 | FEF2 | 20.09 | -155.737 | *Korthalsella* | *sp.* | 07/26/14 |
| K1105 | BIG | 19.688 | -155.269 | *Leptecophylla* | *tameiameiae* | 01/08/15 |
| K1107 | BIG | 19.688 | -155.269 | *Leptecophylla* | *tameiameiae* | 01/08/15 |
| K1117 | BIG | 19.688 | -155.269 | *Leptecophylla* | *tameiameiae* | 01/08/15 |
| K1119 | BIG | 19.688 | -155.269 | *Leptecophylla* | *tameiameiae* | 01/08/15 |
| K1120 | BIG | 19.688 | -155.269 | *Leptecophylla* | *tameiameiae* | 01/08/15 |
| K1201 | BIG | 19.682 | -155.282 | *Leptecophylla* | *tameiameiae* | 01/08/15 |
| K1207 | BIG | 19.682 | -155.282 | *Leptecophylla* | *tameiameiae* | 01/08/15 |
| K1212 | BIG | 19.682 | -155.282 | *Leptecophylla* | *tameiameiae* | 01/08/15 |
| K1216 | BIG | 19.682 | -155.282 | *Leptecophylla* | *tameiameiae* | 01/08/15 |
| K1306 | BIG | 19.68 | -155.298 | *Leptecophylla* | *tameiameiae* | 01/09/15 |
| K1311 | BIG | 19.68 | -155.298 | *Leptecophylla* | *tameiameiae* | 01/09/15 |

| | | | | | | |
|---|---|---|---|---|---|---|
| K1316 | BIG | 19.68 | -155.298 | *Leptecophylla* | *tameiameiae* | 01/09/15 |
| K1320 | BIG | 19.68 | -155.298 | *Leptecophylla* | *tameiameiae* | 01/09/15 |
| K1406 | BIG | 19.677 | -155.314 | *Leptecophylla* | *tameiameiae* | 01/09/15 |
| K1408 | BIG | 19.676 | -155.314 | *Leptecophylla* | *tameiameiae* | 01/09/15 |
| K1411 | BIG | 19.676 | -155.314 | *Leptecophylla* | *tameiameiae* | 01/09/15 |
| K1416 | BIG | 19.677 | -155.314 | *Leptecophylla* | *tameiameiae* | 01/09/15 |
| K1505 | BIG | 19.675 | -155.329 | *Leptecophylla* | *tameiameiae* | 01/09/15 |
| K1508 | BIG | 19.675 | -155.329 | *Leptecophylla* | *tameiameiae* | 01/09/15 |
| K1512 | BIG | 19.675 | -155.33 | *Leptecophylla* | *tameiameiae* | 01/09/15 |
| K1514 | BIG | 19.675 | -155.33 | *Leptecophylla* | *tameiameiae* | 01/09/15 |
| K1518 | BIG | 19.674 | -155.33 | *Leptecophylla* | *tameiameiae* | 01/09/15 |
| K1602 | BIG | 19.671 | -155.346 | *Leptecophylla* | *tameiameiae* | 01/09/15 |
| K1608 | BIG | 19.671 | -155.346 | *Leptecophylla* | *tameiameiae* | 01/09/15 |
| K1610 | BIG | 19.671 | -155.347 | *Leptecophylla* | *tameiameiae* | 01/09/15 |
| K1615 | BIG | 19.671 | -155.347 | *Leptecophylla* | *tameiameiae* | 01/09/15 |
| K1620 | BIG | 19.671 | -155.347 | *Leptecophylla* | *tameiameiae* | 01/09/15 |
| K1803 | BIG | 19.674 | -155.395 | *Leptecophylla* | *tameiameiae* | 01/10/15 |
| K1805 | BIG | 19.674 | -155.395 | *Leptecophylla* | *tameiameiae* | 01/10/15 |
| K1807 | BIG | 19.675 | -155.396 | *Leptecophylla* | *tameiameiae* | 01/10/15 |
| K1809 | BIG | 19.675 | -155.4 | *Leptecophylla* | *tameiameiae* | 01/10/15 |
| K1812 | BIG | 19.674 | -155.401 | *Leptecophylla* | *tameiameiae* | 01/10/15 |
| K1815 | BIG | 19.674 | -155.399 | *Leptecophylla* | *tameiameiae* | 01/10/15 |
| K1904 | BIG | 19.684 | -155.42 | *Leptecophylla* | *tameiameiae* | 01/10/15 |
| K1906 | BIG | 19.684 | -155.42 | *Leptecophylla* | *tameiameiae* | 01/10/15 |
| K1908 | BIG | 19.683 | -155.419 | *Leptecophylla* | *tameiameiae* | 01/10/15 |
| K1913 | BIG | 19.683 | -155.419 | *Leptecophylla* | *tameiameiae* | 01/10/15 |
| K1915 | BIG | 19.683 | -155.419 | *Leptecophylla* | *tameiameiae* | 01/10/15 |
| K2001 | BIG | 19.687 | -155.469 | *Leptecophylla* | *tameiameiae* | 01/10/15 |
| K2007 | BIG | 19.686 | -155.469 | *Leptecophylla* | *tameiameiae* | 01/10/15 |
| K2013 | BIG | 19.685 | -155.467 | *Leptecophylla* | *tameiameiae* | 01/10/15 |
| K2014 | BIG | 19.685 | -155.466 | *Leptecophylla* | *tameiameiae* | 01/10/15 |
| K2015 | BIG | 19.685 | -155.467 | *Leptecophylla* | *tameiameiae* | 01/10/15 |
| O101 | BIG | 19.619 | -155.359 | *Leptecophylla* | *tameiameiae* | 01/11/15 |
| O105 | BIG | 19.619 | -155.359 | *Leptecophylla* | *tameiameiae* | 01/11/15 |
| O112 | BIG | 19.62 | -155.359 | *Leptecophylla* | *tameiameiae* | 01/11/15 |
| O114 | BIG | 19.62 | -155.359 | *Leptecophylla* | *tameiameiae* | 01/11/15 |
| O115 | BIG | 19.62 | -155.359 | *Leptecophylla* | *tameiameiae* | 01/11/15 |
| O203 | BIG | 19.63 | -155.362 | *Leptecophylla* | *tameiameiae* | 01/11/15 |
| O206 | BIG | 19.631 | -155.362 | *Leptecophylla* | *tameiameiae* | 01/11/15 |
| O208 | BIG | 19.631 | -155.362 | *Leptecophylla* | *tameiameiae* | 01/11/15 |
| O213 | BIG | 19.631 | -155.362 | *Leptecophylla* | *tameiameiae* | 01/11/15 |
| O215 | BIG | 19.631 | -155.362 | *Leptecophylla* | *tameiameiae* | 01/11/15 |
| O304 | BIG | 19.64 | -155.364 | *Leptecophylla* | *tameiameiae* | 01/11/15 |
| O308 | BIG | 19.64 | -155.364 | *Leptecophylla* | *tameiameiae* | 01/11/15 |
| O311 | BIG | 19.64 | -155.364 | *Leptecophylla* | *tameiameiae* | 01/11/15 |
| O315 | BIG | 19.641 | -155.364 | *Leptecophylla* | *tameiameiae* | 01/11/15 |
| O316 | BIG | 19.641 | -155.364 | *Leptecophylla* | *tameiameiae* | 01/11/15 |
| O403 | BIG | 19.649 | -155.366 | *Leptecophylla* | *tameiameiae* | 01/11/15 |
| O408 | BIG | 19.649 | -155.366 | *Leptecophylla* | *tameiameiae* | 01/11/15 |
| O417 | BIG | 19.65 | -155.366 | *Leptecophylla* | *tameiameiae* | 01/11/15 |
| O421 | BIG | 19.65 | -155.366 | *Leptecophylla* | *tameiameiae* | 01/11/15 |
| O423 | BIG | 19.65 | -155.366 | *Leptecophylla* | *tameiameiae* | 01/11/15 |
| O503 | BIG | 19.66 | -155.369 | *Leptecophylla* | *tameiameiae* | 01/11/15 |
| O506 | BIG | 19.66 | -155.369 | *Leptecophylla* | *tameiameiae* | 01/11/15 |
| O509 | BIG | 19.661 | -155.369 | *Leptecophylla* | *tameiameiae* | 01/11/15 |
| O512 | BIG | 19.661 | -155.369 | *Leptecophylla* | *tameiameiae* | 01/11/15 |
| O515 | BIG | 19.662 | -155.369 | *Leptecophylla* | *tameiameiae* | 01/11/15 |
| BI024 | FEF2 | 19.412 | -155.251 | *Leptecophylla* | *tameiameiae* | 07/21/14 |
| BI038 | FEF2 | 19.42 | -155.29 | *Leptecophylla* | *tameiameiae* | 07/21/14 |

| | | | | | | |
|---|---|---|---|---|---|---|
| BI044 | FEF2 | 19.474 | -155.358 | *Leptecophylla* | *tameiameiae* | 07/21/14 |
| BI060 | FEF2 | 20.113 | -155.76 | *Leptecophylla* | *tameiameiae* | 07/22/14 |
| BI078 | FEF2 | 19.687 | -155.468 | *Leptecophylla* | *tameiameiae* | 07/23/14 |
| BI095 | FEF2 | 19.676 | -155.329 | *Leptecophylla* | *tameiameiae* | 07/24/14 |
| BI152 | FEF2 | 20.09 | -155.737 | *Leptecophylla* | *tameiameiae* | 07/26/14 |
| K0004 | FEF2 | 22.034 | -159.669 | *Leptecophylla* | *tameiameiae* | 06/15/15 |
| Kan20 | FEF2 | 20.62 | -156.35 | *Leptecophylla* | *tameiameiae* | 06/10/15 |
| SP02 | FEF2 | 21.512 | -158.136 | *Leptecophylla* | *tameiameiae* | 08/28/15 |
| UW02 | FEF2 | 20.773 | -156.235 | *Leptecophylla* | *tameiameiae* | 11/06/14 |
| KEA1 | FEF3 | 21.56 | -158.211 | *Leptecophylla* | *tameiameiae* | 08/18/16 |
| LHL1 | FEF3 | 21.301 | -157.745 | *Leptecophylla* | *tameiameiae* | 08/21/16 |
| MHL1 | FEF3 | 21.308 | -157.745 | *Leptecophylla* | *tameiameiae* | 08/26/16 |
| Mo002 | FEF3 | 21.119 | -156.93 | *Leptecophylla* | *tameiameiae* | 07/18/16 |
| Mo028 | FEF3 | 21.119 | -156.901 | *Leptecophylla* | *tameiameiae* | 07/18/16 |
| Mo054 | FEF3 | 21.1 | -156.914 | *Leptecophylla* | *tameiameiae* | 07/19/16 |
| PUPU2 | FEF3 | 21.64 | -158.018 | *Leptecophylla* | *tameiameiae* | 08/03/16 |
| UW10 | FEF2 | 20.776 | -156.234 | *Lobelia* | *grayana* | 11/06/14 |
| Mo041 | FEF3 | 21.118 | -156.908 | *Lobelia* | *sp.* | 07/18/16 |
| K0068 | FEF2 | 22.15 | -159.615 | *Labordia* | *waialeale* | 06/16/15 |
| KO5 | FEF3 | 21.355 | -157.788 | *Labordia* | *hosakana* | 08/11/16 |
| Mo086 | FEF3 | 21.2 | -157.157 | *Lycium* | *sandwicense* | 07/20/16 |
| Mo097 | FEF3 | 21.201 | -157.158 | *Lycium* | *sandwicense* | 07/20/16 |
| EKH26 | FEF2 | 21.438 | -158.096 | *Lysimachia* | *sp.* | 09/22/15 |
| K0019 | FEF2 | 22.151 | -159.645 | *Lysimachia* | *glutinosa* | 06/15/15 |
| Mo008 | FEF3 | 21.119 | -156.93 | *Lysimachia* | *remyi* | 07/18/16 |
| BI063 | FEF2 | 20.114 | -155.76 | *Melicope* | *cluciifolia* | 07/22/14 |
| BI065 | FEF2 | 20.114 | -155.76 | *Melicope* | *spthulata* | 07/22/14 |
| BI148 | FEF2 | 20.09 | -155.737 | *Melicope* | *cluciifolia* | 07/26/14 |
| BI149 | FEF2 | 20.09 | -155.737 | *Melicope* | *sp.* | 07/26/14 |
| K0014 | FEF2 | 22.138 | -159.653 | *Melicope* | *anisada* | 06/15/15 |
| K0045 | FEF2 | 22.118 | -159.679 | *Melicope* | *barbigera* | 06/15/15 |
| K0108 | FEF2 | 21.975 | -159.507 | *Melicope* | *waialealae* | 06/19/15 |
| wai09B | FEF2 | 20.802 | -156.255 | *Melicope* | *volcanica* | 11/04/14 |
| wai10 | FEF2 | 20.802 | -156.255 | *Melicope* | *cluciifolia* | 11/04/14 |
| wai27 | FEF2 | 20.802 | -156.252 | *Melicope* | *spthulata* | 11/04/14 |
| KO8 | FEF3 | 21.355 | -157.788 | *Melicope* | *clusiifolia* | 08/11/16 |
| L4 | FEF3 | 21.596 | -157.958 | *Melicope* | *clusiifolia* | 07/02/16 |
| Mo036 | FEF3 | 21.119 | -156.899 | *Melicope* | *sp.* | 07/18/16 |
| Mo037 | FEF3 | 21.12 | -156.897 | *Melicope* | *sp.* | 07/18/16 |
| Mo061 | FEF3 | 21.1 | -156.914 | *Melicope* | *sp.* | 07/19/16 |
| K0702 | BIG | 19.693 | -155.202 | *Metrosideros* | *polymorpha* | 01/07/15 |
| K0703 | BIG | 19.693 | -155.202 | *Metrosideros* | *polymorpha* | 01/07/15 |
| K0704 | BIG | 19.693 | -155.202 | *Metrosideros* | *polymorpha* | 01/07/15 |
| K0705 | BIG | 19.694 | -155.202 | *Metrosideros* | *polymorpha* | 01/07/15 |
| K1003 | BIG | 19.695 | -155.251 | *Metrosideros* | *polymorpha* | 01/07/15 |
| K1005 | BIG | 19.695 | -155.252 | *Metrosideros* | *polymorpha* | 01/07/15 |
| K1007 | BIG | 19.695 | -155.252 | *Metrosideros* | *polymorpha* | 01/07/15 |
| K1011 | BIG | 19.696 | -155.251 | *Metrosideros* | *polymorpha* | 01/07/15 |
| K1012 | BIG | 19.696 | -155.251 | *Metrosideros* | *polymorpha* | 01/07/15 |
| K1102 | BIG | 19.688 | -155.269 | *Metrosideros* | *polymorpha* | 01/08/15 |
| K1109 | BIG | 19.688 | -155.269 | *Metrosideros* | *polymorpha* | 01/08/15 |
| K1113 | BIG | 19.687 | -155.268 | *Metrosideros* | *polymorpha* | 01/08/15 |
| K1116 | BIG | 19.688 | -155.268 | *Metrosideros* | *polymorpha* | 01/08/15 |
| K1203 | BIG | 19.682 | -155.282 | *Metrosideros* | *polymorpha* | 01/08/15 |
| K1205 | BIG | 19.682 | -155.282 | *Metrosideros* | *polymorpha* | 01/08/15 |
| K1215 | BIG | 19.682 | -155.282 | *Metrosideros* | *polymorpha* | 01/08/15 |
| K1218 | BIG | 19.682 | -155.282 | *Metrosideros* | *polymorpha* | 01/08/15 |
| K1303 | BIG | 19.681 | -155.298 | *Metrosideros* | *polymorpha* | 01/09/15 |
| K1309 | BIG | 19.68 | -155.298 | *Metrosideros* | *polymorpha* | 01/09/15 |

| | | | | | | |
|---|---|---|---|---|---|---|
| K1313 | BIG | 19.68 | -155.298 | *Metrosideros* | *polymorpha* | 01/09/15 |
| K1318 | BIG | 19.68 | -155.298 | *Metrosideros* | *polymorpha* | 01/09/15 |
| K1404 | BIG | 19.677 | -155.314 | *Metrosideros* | *polymorpha* | 01/09/15 |
| K1407 | BIG | 19.677 | -155.314 | *Metrosideros* | *polymorpha* | 01/09/15 |
| K1410 | BIG | 19.676 | -155.314 | *Metrosideros* | *polymorpha* | 01/09/15 |
| K1415 | BIG | 19.677 | -155.314 | *Metrosideros* | *polymorpha* | 01/09/15 |
| K1419 | BIG | 19.677 | -155.314 | *Metrosideros* | *polymorpha* | 01/09/15 |
| K1503 | BIG | 19.675 | -155.329 | *Metrosideros* | *polymorpha* | 01/09/15 |
| K1507 | BIG | 19.675 | -155.329 | *Metrosideros* | *polymorpha* | 01/09/15 |
| K1510 | BIG | 19.675 | -155.329 | *Metrosideros* | *polymorpha* | 01/09/15 |
| K1516 | BIG | 19.674 | -155.33 | *Metrosideros* | *polymorpha* | 01/09/15 |
| K1519 | BIG | 19.674 | -155.331 | *Metrosideros* | *polymorpha* | 01/09/15 |
| K1603 | BIG | 19.671 | -155.346 | *Metrosideros* | *polymorpha* | 01/09/15 |
| K1605 | BIG | 19.671 | -155.346 | *Metrosideros* | *polymorpha* | 01/09/15 |
| K1612 | BIG | 19.67 | -155.346 | *Metrosideros* | *polymorpha* | 01/09/15 |
| K1617 | BIG | 19.671 | -155.347 | *Metrosideros* | *polymorpha* | 01/09/15 |
| K1619 | BIG | 19.671 | -155.347 | *Metrosideros* | *polymorpha* | 01/09/15 |
| K1806 | BIG | 19.675 | -155.395 | *Metrosideros* | *polymorpha* | 01/10/15 |
| K1808 | BIG | 19.675 | -155.4 | *Metrosideros* | *polymorpha* | 01/10/15 |
| K1811 | BIG | 19.674 | -155.401 | *Metrosideros* | *polymorpha* | 01/10/15 |
| K1813 | BIG | 19.674 | -155.399 | *Metrosideros* | *polymorpha* | 01/10/15 |
| K1902 | BIG | 19.684 | -155.42 | *Metrosideros* | *polymorpha* | 01/10/15 |
| K1905 | BIG | 19.684 | -155.419 | *Metrosideros* | *polymorpha* | 01/10/15 |
| K1909 | BIG | 19.683 | -155.419 | *Metrosideros* | *polymorpha* | 01/10/15 |
| K1911 | BIG | 19.683 | -155.419 | *Metrosideros* | *polymorpha* | 01/10/15 |
| K1914 | BIG | 19.683 | -155.419 | *Metrosideros* | *polymorpha* | 01/10/15 |
| K2003 | BIG | 19.687 | -155.469 | *Metrosideros* | *polymorpha* | 01/10/15 |
| K2005 | BIG | 19.687 | -155.469 | *Metrosideros* | *polymorpha* | 01/10/15 |
| K2009 | BIG | 19.686 | -155.468 | *Metrosideros* | *polymorpha* | 01/10/15 |
| K2010 | BIG | 19.685 | -155.467 | *Metrosideros* | *polymorpha* | 01/10/15 |
| K2012 | BIG | 19.685 | -155.467 | *Metrosideros* | *polymorpha* | 01/10/15 |
| O104 | BIG | 19.619 | -155.359 | *Metrosideros* | *polymorpha* | 01/11/15 |
| O107 | BIG | 19.619 | -155.359 | *Metrosideros* | *polymorpha* | 01/11/15 |
| O109 | BIG | 19.619 | -155.359 | *Metrosideros* | *polymorpha* | 01/11/15 |
| O111 | BIG | 19.62 | -155.359 | *Metrosideros* | *polymorpha* | 01/11/15 |
| O116 | BIG | 19.62 | -155.359 | *Metrosideros* | *polymorpha* | 01/11/15 |
| O202 | BIG | 19.63 | -155.362 | *Metrosideros* | *polymorpha* | 01/11/15 |
| O205 | BIG | 19.631 | -155.362 | *Metrosideros* | *polymorpha* | 01/11/15 |
| O209 | BIG | 19.631 | -155.362 | *Metrosideros* | *polymorpha* | 01/11/15 |
| O211 | BIG | 19.631 | -155.362 | *Metrosideros* | *polymorpha* | 01/11/15 |
| O214 | BIG | 19.631 | -155.362 | *Metrosideros* | *polymorpha* | 01/11/15 |
| O305 | BIG | 19.64 | -155.364 | *Metrosideros* | *polymorpha* | 01/11/15 |
| O306 | BIG | 19.64 | -155.364 | *Metrosideros* | *polymorpha* | 01/11/15 |
| O309 | BIG | 19.64 | -155.364 | *Metrosideros* | *polymorpha* | 01/11/15 |
| O312 | BIG | 19.64 | -155.364 | *Metrosideros* | *polymorpha* | 01/11/15 |
| O314 | BIG | 19.641 | -155.364 | *Metrosideros* | *polymorpha* | 01/11/15 |
| O410 | BIG | 19.649 | -155.366 | *Metrosideros* | *polymorpha* | 01/11/15 |
| O412 | BIG | 19.65 | -155.366 | *Metrosideros* | *polymorpha* | 01/11/15 |
| O418 | BIG | 19.65 | -155.366 | *Metrosideros* | *polymorpha* | 01/11/15 |
| O422 | BIG | 19.65 | -155.366 | *Metrosideros* | *polymorpha* | 01/11/15 |
| O502 | BIG | 19.66 | -155.369 | *Metrosideros* | *polymorpha* | 01/11/15 |
| O505 | BIG | 19.66 | -155.369 | *Metrosideros* | *polymorpha* | 01/11/15 |
| O511 | BIG | 19.661 | -155.369 | *Metrosideros* | *polymorpha* | 01/11/15 |
| O514 | BIG | 19.662 | -155.369 | *Metrosideros* | *polymorpha* | 01/11/15 |
| X69cv08 | FEF2 | 21.515 | -158.161 | *Metrosideros* | *polymorpha* | 09/08/15 |
| BI010 | FEF2 | 19.415 | -155.237 | *Metrosideros* | *polymorpha* | 07/21/14 |
| BI025 | FEF2 | 19.412 | -155.251 | *Metrosideros* | *polymorpha* | 07/21/14 |
| BI041A | FEF2 | 19.42 | -155.29 | *Metrosideros* | *polymorpha* | 07/21/14 |
| BI041B | FEF2 | 19.42 | -155.29 | *Metrosideros* | *polymorpha* | 07/21/14 |

| | | | | | | |
|---|---|---|---|---|---|---|
| BI076 | FEF2 | 19.687 | -155.466 | *Metrosideros* | *polymorpha* | 07/23/14 |
| BI086 | FEF2 | 19.417 | -154.951 | *Metrosideros* | *polymorpha* | 07/24/14 |
| BI112 | FEF2 | 19.672 | -155.338 | *Metrosideros* | *polymorpha* | 07/24/14 |
| BI118 | FEF2 | 19.676 | -155.384 | *Metrosideros* | *polymorpha* | 07/25/14 |
| BI136 | FEF2 | 19.649 | -155.372 | *Metrosideros* | *polymorpha* | 07/25/14 |
| BI157 | FEF2 | 20.09 | -155.737 | *Metrosideros* | *polymorpha* | 07/26/14 |
| BI213 | FEF2 | 19.616 | -155.928 | *Metrosideros* | *polymorpha* | 03/25/15 |
| BI233 | FEF2 | 19.615 | -155.931 | *Metrosideros* | *polymorpha* | 03/25/15 |
| BI244 | FEF2 | 19.115 | -155.82 | *Metrosideros* | *polymorpha* | 03/25/15 |
| EKH24 | FEF2 | 21.438 | -158.096 | *Metrosideros* | *polymorpha* | 09/22/15 |
| K0020 | FEF2 | 22.151 | -159.645 | *Metrosideros* | *polymorpha* | 06/15/15 |
| K0043B | FEF2 | 22.118 | -159.679 | *Metrosideros* | *polymorpha* | 06/15/15 |
| K0049 | FEF2 | 22.148 | -159.629 | *Metrosideros* | *polymorpha* | 06/16/15 |
| K0122A | FEF2 | 22.097 | -159.745 | *Metrosideros* | *polymorpha* | 06/20/15 |
| UW05 | FEF2 | 20.773 | -156.235 | *Metrosideros* | *polymorpha* | 11/06/14 |
| UW08 | FEF2 | 20.773 | -156.236 | *Metrosideros* | *polymorpha* | 11/06/14 |
| HL8 | FEF3 | 21.324 | -157.742 | *Metrosideros* | *rugosa* | 07/21/16 |
| KO4 | FEF3 | 21.355 | -157.788 | *Metrosideros* | *polymorpha* | 08/11/16 |
| KO7 | FEF3 | 21.355 | -157.788 | *Metrosideros* | *rugosa* | 08/11/16 |
| MC13 | FEF3 | 21.333 | -157.81 | *Metrosideros* | *tremuloides* | 07/15/16 |
| MC4 | FEF3 | 21.336 | -157.81 | *Metrosideros* | *tremuloides* | 07/15/16 |
| MHL3 | FEF3 | 21.308 | -157.745 | *Metrosideros* | *polymorpha* | 08/26/16 |
| Mo020 | FEF3 | 21.118 | -156.905 | *Metrosideros* | *polymorpha* | 07/18/16 |
| Mo027 | FEF3 | 21.119 | -156.901 | *Metrosideros* | *polymorpha* | 07/18/16 |
| Mo055 | FEF3 | 21.1 | -156.914 | *Metrosideros* | *polymorpha* | 07/19/16 |
| POA1 | FEF3 | 21.53 | -157.941 | *Metrosideros* | *polymorpha* | 10/02/16 |
| POA19 | FEF3 | 21.534 | -157.924 | *Metrosideros* | *polymorpha* | 10/02/16 |
| POAR3 | FEF3 | 21.526 | -157.918 | *Metrosideros* | *polymorpha* | 10/02/16 |
| BI075 | FEF2 | 19.686 | -155.467 | *Myoporum* | *sandwicense* | 07/23/14 |
| BI139 | FEF2 | 19.649 | -155.372 | *Myoporum* | *sandwicense* | 07/25/14 |
| Kan08 | FEF2 | 20.61 | -156.34 | *Myoporum* | *sandwicensis* | 06/10/15 |
| Maka10 | FEF2 | 21.293 | -157.661 | *Myoporum* | *sandwicense* | 07/21/15 |
| Mo045 | FEF3 | 21.091 | -156.928 | *Myoporum* | *sandwicense* | 07/19/16 |
| BI020 | FEF2 | 19.412 | -155.251 | *Myrsine* | *umbellata* | 07/21/14 |
| BI057 | FEF2 | 20.113 | -155.759 | *Myrsine* | *lessertiana* | 07/22/14 |
| BI071 | FEF2 | 19.687 | -155.466 | *Myrsine* | *sp.* | 07/23/14 |
| BI103 | FEF2 | 19.674 | -155.33 | *Myrsine* | *sp.* | 07/24/14 |
| BI109 | FEF2 | 19.672 | -155.338 | *Myrsine* | *sp.* | 07/24/14 |
| BI128 | FEF2 | 19.662 | -155.387 | *Myrsine* | *lessertiana* | 07/25/14 |
| BI140 | FEF2 | 19.649 | -155.372 | *Myrsine* | *lessertiana* | 07/25/14 |
| BI150 | FEF2 | 20.09 | -155.737 | *Myrsine* | *sandwicensis* | 07/26/14 |
| BI161 | FEF2 | 20.091 | -155.738 | *Myrsine* | *sandwicensis* | 07/26/14 |
| EKH17 | FEF2 | 21.439 | -158.097 | *Myrsine* | *lessertiana* | 09/22/15 |
| K0017 | FEF2 | 22.138 | -159.653 | *Myrsine* | *alixopholia* | 06/15/15 |
| K0060 | FEF2 | 22.154 | -159.619 | *Myrsine* | *sp.* | 06/16/15 |
| K0103 | FEF2 | 21.975 | -159.508 | *Myrsine* | *helleri* | 06/19/15 |
| KO10 | FEF3 | 21.354 | -157.788 | *Myrsine* | *sandwicensis* | 08/11/16 |
| Mo013 | FEF3 | 21.117 | -156.919 | *Myrsine* | *lessertiana* | 07/18/16 |
| Maka09 | FEF2 | 21.292 | -157.662 | *Nama* | *sandwicensis* | 07/21/15 |
| K0021A | FEF2 | 22.151 | -159.644 | *Nertera* | *granadensis* | 06/15/15 |
| Mo017 | FEF3 | 21.118 | -156.905 | *Nertera* | *granadensis* | 07/18/16 |
| Mo077 | FEF3 | 21.099 | -156.914 | *Nertera* | *granadensis* | 07/19/16 |
| POAR1 | FEF3 | 21.526 | -157.918 | *Nertera* | *granadensis* | 10/02/16 |
| X69cv09 | FEF2 | 21.515 | -158.161 | *Nestegis* | *sandwicensis* | 09/08/15 |
| BI207 | FEF2 | 19.615 | -155.927 | *Nestegis* | *sandwicensis* | 03/25/15 |
| BI231 | FEF2 | 19.615 | -155.931 | *Nestegis* | *sandwiciensis* | 03/25/15 |
| BI243 | FEF2 | 19.115 | -155.82 | *Nestegis* | *sandwicensis* | 03/25/15 |
| EKH15 | FEF2 | 21.439 | -158.096 | *Nestegis* | *sandwicensis* | 09/22/15 |
| EKH4 | FEF2 | 21.439 | -158.095 | *Nestegis* | *sandwicensis* | 09/22/15 |

| | | | | | | |
|---|---|---|---|---|---|---|
| EKH6 | FEF2 | 21.439 | -158.095 | *Nestegis* | *sandwicensis* | 09/22/15 |
| K0013 | FEF2 | 22.138 | -159.653 | *Nestegis* | *sandwicensis* | 06/15/15 |
| K0035B | FEF2 | 22.148 | -159.636 | *Nestegis* | *sandwicensis* | 06/15/15 |
| Kap20 | FEF2 | 20.933 | -156.63 | *Nestegis* | *sandwicensis* | 06/11/15 |
| Mo066 | FEF3 | 21.102 | -156.909 | *Nestegis* | *sandwicensis* | 07/19/16 |
| BI237 | FEF2 | 19.111 | -155.824 | *Osteomeles* | *anthyllidifolia* | 03/25/15 |
| Kap2P7 | FEF2 | 20.935 | -156.64 | *Osteomeles* | *anthyllidifolia* | 06/11/15 |
| LHL3 | FEF3 | 21.301 | -157.745 | *Osteomeles* | *anthyllidifolia* | 08/21/16 |
| Mo056 | FEF3 | 21.1 | -156.914 | *Osteomeles* | *anthyllidifolia* | 07/19/16 |
| BI083 | FEF2 | 19.417 | -154.951 | *Perrottetia* | *sandwicensis* | 07/24/14 |
| BI113 | FEF2 | 19.672 | -155.338 | *Perrottetia* | *sandwicensis* | 07/24/14 |
| EKH18 | FEF2 | 21.439 | -158.097 | *Perrottetia* | *sandwicensis* | 09/22/15 |
| K0037 | FEF2 | 22.148 | -159.636 | *Perrottetia* | *sandwicensis* | 06/15/15 |
| SP21 | FEF2 | 21.512 | -158.137 | *Perrottetia* | *sandwicensis* | 08/28/15 |
| POAR7 | FEF3 | 21.527 | -157.92 | *Phyllostegia* | *grandiflora* | 10/02/16 |
| wai12 | FEF2 | 20.802 | -156.255 | *Phytolacca* | *sandwicensis* | 11/04/14 |
| BI141 | FEF2 | 19.648 | -155.372 | *Pipturus* | *albidus* | 07/25/14 |
| BI165 | FEF2 | 20.091 | -155.739 | *Pipturus* | *albidus* | 07/26/14 |
| BI211 | FEF2 | 19.615 | -155.927 | *Pipturus* | *albidus* | 03/25/15 |
| BI220 | FEF2 | 19.615 | -155.932 | *Pipturus* | *albidus* | 03/25/15 |
| BI241 | FEF2 | 19.114 | -155.822 | *Pipturus* | *albidus* | 03/25/15 |
| EKH12 | FEF2 | 21.439 | -158.096 | *Pipturus* | *albidus* | 09/22/15 |
| MC2 | FEF3 | 21.337 | -157.811 | *Pipturus* | *albidus* | 07/15/16 |
| BI236 | FEF2 | 19.615 | -155.931 | *Pisonia* | *umbellifera* | 03/25/15 |
| BI242 | FEF2 | 19.115 | -155.82 | *Pisonia* | *sandwicensis* | 03/25/15 |
| EKH10 | FEF2 | 21.439 | -158.096 | *Pisonia* | *sandwicensis* | 09/22/15 |
| K0080 | FEF2 | 22.213 | -159.577 | *Pisonia* | *umbellifera* | 06/18/15 |
| JWA323 | FEF3 | 21.336 | -157.81 | *Pisonia* | *sp.* | 07/15/16 |
| BI228 | FEF2 | 19.616 | -155.931 | *Pittosporum* | *hosmori* | 03/25/15 |
| EKH9 | FEF2 | 21.439 | -158.095 | *Pittosporum* | *sp.* | 09/22/15 |
| K0057 | FEF2 | 22.15 | -159.623 | *Pittosporum* | *kauaiense* | 06/16/15 |
| K0113 | FEF2 | 21.981 | -159.502 | *Pittosporum* | *glabrum* | 06/19/15 |
| KB01 | FEF2 | 21.507 | -158.144 | *Pittosporum* | *sp.* | 06/11/15 |
| KB09 | FEF2 | 21.507 | -158.144 | *Pittosporum* | *sp.* | 06/11/15 |
| KB10 | FEF2 | 21.507 | -158.144 | *Pittosporum* | *sp.* | 06/11/15 |
| KB12 | FEF2 | 21.504 | -158.147 | *Pittosporum* | *sp.* | 06/11/15 |
| KB14 | FEF2 | 21.503 | -158.148 | *Pittosporum* | *sp.* | 06/11/15 |
| Mo025 | FEF3 | 21.118 | -156.903 | *Pittosporum* | *glabrum* | 07/18/16 |
| Mo064 | FEF3 | 21.102 | -156.911 | *Pittosporum* | *glabrum* | 07/19/16 |
| K0016 | FEF2 | 22.138 | -159.653 | *Planchonella* | *sandwicensis* | 06/15/15 |
| MC10 | FEF3 | 21.334 | -157.81 | *Planchonella* | *sandwicensis* | 07/15/16 |
| Kap13 | FEF2 | 20.933 | -156.632 | *Plectranthus* | *parvifolia* | 06/11/15 |
| BI210 | FEF2 | 19.615 | -155.927 | *Polyscias* | *hawaiiensis* | 03/25/15 |
| K0033 | FEF2 | 22.149 | -159.641 | *Polyscias* | *sp.* | 06/15/15 |
| K0120 | FEF2 | 21.984 | -159.501 | *Polyscias* | *waialealae* | 06/19/15 |
| Kan07 | FEF2 | 20.61 | -156.34 | *Polyscias* | *sandwicensis* | 06/10/15 |
| HL6 | FEF3 | 21.322 | -157.742 | *Polyscias* | *oahuensis* | 07/21/16 |
| POA18 | FEF3 | 21.534 | -157.924 | *Polyscias* | *oahuensis* | 10/02/16 |
| Kan17 | FEF2 | 20.62 | -156.35 | *Pouteria* | *spathulata* | 06/10/15 |
| BI223 | FEF2 | 19.615 | -155.932 | *Psychotria* | *hawaiiensis* | 03/25/15 |
| EKH8 | FEF2 | 21.439 | -158.095 | *Psychotria* | *sp.* | 09/22/15 |
| K0010 | FEF2 | 22.138 | -159.653 | *Psychotria* | *sp.* | 06/15/15 |
| K0011 | FEF2 | 22.138 | -159.653 | *Psychotria* | *sp.* | 06/15/15 |
| K0081 | FEF2 | 22.212 | -159.576 | *Psychotria* | *mariniana* | 06/18/15 |
| K0110 | FEF2 | 21.977 | -159.507 | *Psychotria* | *hexandra* | 06/19/15 |
| KO16 | FEF3 | 21.352 | -157.793 | *Psychotria* | *mariniana* | 08/11/16 |
| MC11 | FEF3 | 21.334 | -157.81 | *Psychotria* | *mariniana* | 07/15/16 |
| MC3 | FEF3 | 21.337 | -157.811 | *Psychotria* | *kaduana* | 07/15/16 |
| Mo014 | FEF3 | 21.117 | -156.919 | *Psychotria* | *sp.* | 07/18/16 |

| | | | | | | |
|---|---|---|---|---|---|---|
| POA22 | FEF3 | 21.533 | -157.925 | *Psychotria* | *mariniana* | 10/02/16 |
| BI089 | FEF2 | 19.448 | -154.862 | *Psydrax* | *odorata* | 07/24/14 |
| BI203 | FEF2 | 19.615 | -155.927 | *Psydrax* | *odorata* | 03/25/15 |
| BI232 | FEF2 | 19.615 | -155.931 | *Psydrax* | *odorata* | 03/25/15 |
| BI238 | FEF2 | 19.111 | -155.824 | *Psydrax* | *odorata* | 03/25/15 |
| K0082 | FEF2 | 22.212 | -159.576 | *Psydrax* | *odorata* | 06/18/15 |
| Kan14 | FEF2 | 20.61 | -156.35 | *Rauvolfia* | *sandwicensis* | 06/10/15 |
| Kap14 | FEF2 | 20.933 | -156.631 | *Rauvolfia* | *sandwicensis* | 06/11/15 |
| BI061 | FEF2 | 20.113 | -155.76 | *Rubus* | *hawaiensis* | 07/22/14 |
| BI125 | FEF2 | 19.665 | -155.388 | *Rubus* | *hawaiensis* | 07/25/14 |
| BI156 | FEF2 | 20.09 | -155.737 | *Rubus* | *hawaiensis* | 07/26/14 |
| UW13 | FEF2 | 20.776 | -156.234 | *Rubus* | *hawaiensis* | 11/06/14 |
| BI067 | FEF2 | 19.687 | -155.466 | *Santalum* | *paniculatum* | 07/23/14 |
| BI131 | FEF2 | 19.652 | -155.376 | *Santalum* | *paniculatum* | 07/25/14 |
| BI201 | FEF2 | 19.485 | -155.269 | *Santalum* | *sp.* | 03/24/15 |
| K0043A | FEF2 | 22.118 | -159.679 | *Santalum* | *sp.* | 06/15/15 |
| Kan19 | FEF2 | 20.62 | -156.35 | *Santalum* | *ellipticum* | 06/10/15 |
| Maka20 | FEF2 | 21.314 | -157.661 | *Santalum* | *ellipticum* | 07/21/15 |
| HL3 | FEF3 | 21.315 | -157.743 | *Santalum* | *freycinetianum* | 07/21/16 |
| KEA6 | FEF3 | 21.566 | -158.21 | *Santalum* | *elipticum* | 08/18/16 |
| BI092 | FEF2 | 19.445 | -154.856 | *Scaevola* | *sp.* | 07/24/14 |
| BI186A | FEF2 | 19.339 | -155.274 | *Scaevola* | *kilaueae* | 03/22/15 |
| BI186B | FEF2 | 19.407 | -155.253 | *Scaevola* | *kilaueae* | 03/21/15 |
| BI190 | FEF2 | 19.448 | -155.202 | *Scaevola* | *chamissoniana* | 03/24/15 |
| BI226 | FEF2 | 19.616 | -155.931 | *Scaevola* | *chamissoniana* | 03/25/15 |
| EKH28 | FEF2 | 21.438 | -158.096 | *Scaevola* | *mollis* | 09/22/15 |
| K0006 | FEF2 | 22.034 | -159.669 | *Scaevola* | *guadichaudii* | 06/15/15 |
| K0007 | FEF2 | 22.034 | -159.669 | *Scaevola* | *guadichaudii* | 06/15/15 |
| K0009 | FEF2 | 22.138 | -159.653 | *Scaevola* | *procera* | 06/15/15 |
| K0021B | FEF2 | 22.151 | -159.644 | *Scaevola* | *procera* | 06/15/15 |
| K0036 | FEF2 | 22.148 | -159.636 | *Scaevola* | *glabra* | 06/15/15 |
| K0048 | FEF2 | 22.148 | -159.629 | *Scaevola* | *procera* | 06/16/15 |
| K0055 | FEF2 | 22.15 | -159.623 | *Scaevola* | *glabra* | 06/16/15 |
| K0078 | FEF2 | 21.893 | -159.405 | *Scaevola* | *taccada* | 06/17/15 |
| K0115 | FEF2 | 21.983 | -159.501 | *Scaevola* | *mollis* | 06/19/15 |
| Maka13 | FEF2 | 21.316 | -157.663 | *Scaevola* | *sericea* | 07/21/15 |
| KO14 | FEF3 | 21.353 | -157.789 | *Scaevola* | *mollis* | 08/11/16 |
| Mo010 | FEF3 | 21.119 | -156.926 | *Scaevola* | *mollis* | 07/18/16 |
| Mo015 | FEF3 | 21.117 | -156.919 | *Scaevola* | *mollis* | 07/18/16 |
| Mo042 | FEF3 | 21.123 | -156.917 | *Scaevola* | *chamissoniana* | 07/18/16 |
| Mo063 | FEF3 | 21.101 | -156.913 | *Scaevola* | *gaudichaudii* | 07/19/16 |
| Mo068 | FEF3 | 21.102 | -156.909 | *Scaevola* | *chamissoniana* | 07/19/16 |
| Mo078 | FEF3 | 21.099 | -156.914 | *Scaevola* | *gaudichaudii* | 07/19/16 |
| Mo084 | FEF3 | 21.2 | -157.157 | *Scaevola* | *taccada* | 07/20/16 |
| Mo089 | FEF3 | 21.2 | -157.157 | *Scaevola* | *taccada* | 07/20/16 |
| Mo093 | FEF3 | 21.201 | -157.158 | *Scaevola* | *taccada* | 07/20/16 |
| NS2 | FEF3 | 21.58 | -158.214 | *Scaevola* | *taccada* | 08/03/16 |
| NS6 | FEF3 | 21.581 | -158.207 | *Scaevola* | *taccada* | 08/03/16 |
| NS7 | FEF3 | 21.58 | -158.165 | *Scaevola* | *taccada* | 08/03/16 |
| POAR11 | FEF3 | 21.532 | -157.921 | *Scaevola* | *mollis* | 10/02/16 |
| EKH23 | FEF2 | 21.438 | -158.096 | *Schinus* | *terebinthifolia* | 09/22/15 |
| Maka07 | FEF2 | 21.292 | -157.662 | *Sesbania* | *tomentosa* | 07/21/15 |
| Mo088 | FEF3 | 21.2 | -157.157 | *Sesbania* | *tomentosa* | 07/20/16 |
| BI007 | FEF2 | 19.414 | -155.238 | *Sesuvium* | *portulacastrum* | 07/21/14 |
| K0071 | FEF2 | 21.893 | -159.41 | *Sesuvium* | *portulacastrum* | 06/17/15 |
| Maka19 | FEF2 | 21.315 | -157.661 | *Sesuvium* | *portulacastrum* | 07/21/15 |
| Mo090 | FEF3 | 21.2 | -157.157 | *Sesuvium* | *portulacastrum* | 07/20/16 |
| BI072 | FEF2 | 19.687 | -155.467 | *Sicyos* | *lanceoloidea* | 07/23/14 |
| Kap01 | FEF2 | 20.934 | -156.64 | *Sida* | *rotundifolia* | 06/11/15 |

| | | | | | | |
|---|---|---|---|---|---|---|
| Maka05 | FEF2 | 21.289 | -157.667 | *Sida* | *fallax* | 07/21/15 |
| MHL5 | FEF3 | 21.307 | -157.745 | *Sida* | *fallax* | 08/26/16 |
| Mo046 | FEF3 | 21.091 | -156.928 | *Sida* | *fallax* | 07/19/16 |
| Mo091 | FEF3 | 21.2 | -157.157 | *Sida* | *fallax* | 07/20/16 |
| Mo085 | FEF3 | 21.2 | -157.157 | *Solanum* | *nelsonii* | 07/20/16 |
| BI048 | FEF2 | 19.458 | -155.34 | *Sophora* | *chrysophylla* | 07/21/14 |
| BI077 | FEF2 | 19.687 | -155.468 | *Sophora* | *chrysophylla* | 07/23/14 |
| UW15 | FEF2 | 20.776 | -156.234 | *Sophora* | *chrysophylla* | 11/06/14 |
| wai25 | FEF2 | 20.8 | -156.252 | *Sophora* | *chrysophylla* | 11/04/14 |
| Mo080 | FEF3 | 21.099 | -156.914 | *Sophora* | *chrysophylla* | 07/19/16 |
| K0067 | FEF2 | 22.152 | -159.615 | *Stenogyne* | *purpurea* | 06/16/15 |
| Mo038 | FEF3 | 21.12 | -156.897 | *Stenogyne* | *kamehamehae* | 07/18/16 |
| X69cv03 | FEF2 | 21.515 | -158.161 | *Styphelia* | *tameiameiae* | 09/08/15 |
| KO1 | FEF3 | 21.355 | -157.788 | *Syzygium* | *sandwicensis* | 08/11/16 |
| BI059 | FEF2 | 20.113 | -155.759 | *Trematolobelia* | *sp.* | 07/22/14 |
| K0063A | FEF2 | 22.153 | -159.617 | *Trematolobelia* | *kauaiensis* | 06/16/15 |
| K0063B | FEF2 | 22.153 | -159.617 | *Trematolobelia* | *kauaiensis* | 06/16/15 |
| POAR5 | FEF3 | 21.526 | -157.918 | *Trematolobelia* | *macrostachys* | 10/02/16 |
| K0086 | FEF2 | 22.211 | -159.578 | *Urera* | *glabra* | 06/18/15 |
| K0090 | FEF2 | 22.209 | -159.579 | *Touchardia* | *latifolia* | 06/18/15 |
| MC7 | FEF3 | 21.336 | -157.811 | *Touchardia* | *latifolia* | 07/15/16 |
| K1001 | BIG | 19.695 | -155.25 | *Vaccinium* | *reticulatum* | 01/07/15 |
| K1002 | BIG | 19.695 | -155.251 | *Vaccinium* | *reticulatum* | 01/07/15 |
| K1004 | BIG | 19.695 | -155.252 | *Vaccinium* | *reticulatum* | 01/07/15 |
| K1006 | BIG | 19.695 | -155.252 | *Vaccinium* | *reticulatum* | 01/07/15 |
| K1008 | BIG | 19.695 | -155.252 | *Vaccinium* | *reticulatum* | 01/07/15 |
| K1009 | BIG | 19.695 | -155.251 | *Vaccinium* | *reticulatum* | 01/07/15 |
| K1010 | BIG | 19.696 | -155.251 | *Vaccinium* | *reticulatum* | 01/07/15 |
| K1101 | BIG | 19.688 | -155.269 | *Vaccinium* | *reticulatum* | 01/08/15 |
| K1103 | BIG | 19.688 | -155.269 | *Vaccinium* | *reticulatum* | 01/08/15 |
| K1104 | BIG | 19.688 | -155.269 | *Vaccinium* | *reticulatum* | 01/08/15 |
| K1108 | BIG | 19.688 | -155.269 | *Vaccinium* | *reticulatum* | 01/08/15 |
| K1112 | BIG | 19.687 | -155.268 | *Vaccinium* | *reticulatum* | 01/08/15 |
| K1114 | BIG | 19.688 | -155.268 | *Vaccinium* | *reticulatum* | 01/08/15 |
| K1115 | BIG | 19.688 | -155.268 | *Vaccinium* | *reticulatum* | 01/08/15 |
| K1118 | BIG | 19.688 | -155.269 | *Vaccinium* | *reticulatum* | 01/08/15 |
| K1202 | BIG | 19.682 | -155.282 | *Vaccinium* | *reticulatum* | 01/08/15 |
| K1204 | BIG | 19.682 | -155.282 | *Vaccinium* | *reticulatum* | 01/08/15 |
| K1206 | BIG | 19.682 | -155.282 | *Vaccinium* | *reticulatum* | 01/08/15 |
| K1208 | BIG | 19.682 | -155.282 | *Vaccinium* | *reticulatum* | 01/08/15 |
| K1209 | BIG | 19.682 | -155.282 | *Vaccinium* | *reticulatum* | 01/08/15 |
| K1211 | BIG | 19.682 | -155.282 | *Vaccinium* | *reticulatum* | 01/08/15 |
| K1213 | BIG | 19.682 | -155.282 | *Vaccinium* | *reticulatum* | 01/08/15 |
| K1214 | BIG | 19.682 | -155.282 | *Vaccinium* | *reticulatum* | 01/08/15 |
| K1217 | BIG | 19.682 | -155.282 | *Vaccinium* | *reticulatum* | 01/08/15 |
| K1302 | BIG | 19.681 | -155.298 | *Vaccinium* | *reticulatum* | 01/09/15 |
| K1304 | BIG | 19.681 | -155.298 | *Vaccinium* | *reticulatum* | 01/09/15 |
| K1305 | BIG | 19.68 | -155.298 | *Vaccinium* | *reticulatum* | 01/09/15 |
| K1307 | BIG | 19.68 | -155.298 | *Vaccinium* | *reticulatum* | 01/09/15 |
| K1308 | BIG | 19.68 | -155.298 | *Vaccinium* | *reticulatum* | 01/09/15 |
| K1310 | BIG | 19.68 | -155.298 | *Vaccinium* | *reticulatum* | 01/09/15 |
| K1312 | BIG | 19.68 | -155.298 | *Vaccinium* | *reticulatum* | 01/09/15 |
| K1314 | BIG | 19.68 | -155.298 | *Vaccinium* | *reticulatum* | 01/09/15 |
| K1315 | BIG | 19.68 | -155.298 | *Vaccinium* | *reticulatum* | 01/09/15 |
| K1317 | BIG | 19.68 | -155.298 | *Vaccinium* | *reticulatum* | 01/09/15 |
| K1319 | BIG | 19.68 | -155.298 | *Vaccinium* | *reticulatum* | 01/09/15 |
| K1402 | BIG | 19.677 | -155.314 | *Vaccinium* | *reticulatum* | 01/09/15 |
| K1403 | BIG | 19.677 | -155.314 | *Vaccinium* | *reticulatum* | 01/09/15 |
| K1405 | BIG | 19.677 | -155.314 | *Vaccinium* | *reticulatum* | 01/09/15 |

| | | | | | | |
|---|---|---|---|---|---|---|
| K1409 | BIG | 19.676 | -155.314 | *Vaccinium* | *reticulatum* | 01/09/15 |
| K1412 | BIG | 19.676 | -155.314 | *Vaccinium* | *reticulatum* | 01/09/15 |
| K1413 | BIG | 19.677 | -155.314 | *Vaccinium* | *reticulatum* | 01/09/15 |
| K1414 | BIG | 19.677 | -155.314 | *Vaccinium* | *reticulatum* | 01/09/15 |
| K1417 | BIG | 19.677 | -155.314 | *Vaccinium* | *reticulatum* | 01/09/15 |
| K1418 | BIG | 19.677 | -155.314 | *Vaccinium* | *reticulatum* | 01/09/15 |
| K1501 | BIG | 19.675 | -155.329 | *Vaccinium* | *reticulatum* | 01/09/15 |
| K1502 | BIG | 19.675 | -155.329 | *Vaccinium* | *reticulatum* | 01/09/15 |
| K1504 | BIG | 19.675 | -155.329 | *Vaccinium* | *reticulatum* | 01/09/15 |
| K1506 | BIG | 19.675 | -155.329 | *Vaccinium* | *reticulatum* | 01/09/15 |
| K1509 | BIG | 19.675 | -155.329 | *Vaccinium* | *reticulatum* | 01/09/15 |
| K1511 | BIG | 19.675 | -155.329 | *Vaccinium* | *reticulatum* | 01/09/15 |
| K1513 | BIG | 19.674 | -155.33 | *Vaccinium* | *reticulatum* | 01/09/15 |
| K1515 | BIG | 19.674 | -155.33 | *Vaccinium* | *reticulatum* | 01/09/15 |
| K1517 | BIG | 19.674 | -155.33 | *Vaccinium* | *reticulatum* | 01/09/15 |
| K1601 | BIG | 19.671 | -155.346 | *Vaccinium* | *reticulatum* | 01/09/15 |
| K1606 | BIG | 19.671 | -155.346 | *Vaccinium* | *reticulatum* | 01/09/15 |
| K1607 | BIG | 19.671 | -155.346 | *Vaccinium* | *reticulatum* | 01/09/15 |
| K1609 | BIG | 19.671 | -155.347 | *Vaccinium* | *reticulatum* | 01/09/15 |
| K1611 | BIG | 19.671 | -155.347 | *Vaccinium* | *reticulatum* | 01/09/15 |
| K1613 | BIG | 19.671 | -155.347 | *Vaccinium* | *reticulatum* | 01/09/15 |
| K1614 | BIG | 19.671 | -155.347 | *Vaccinium* | *reticulatum* | 01/09/15 |
| K1616 | BIG | 19.671 | -155.347 | *Vaccinium* | *reticulatum* | 01/09/15 |
| K1618 | BIG | 19.671 | -155.347 | *Vaccinium* | *reticulatum* | 01/09/15 |
| K1801 | BIG | 19.674 | -155.395 | *Vaccinium* | *reticulatum* | 01/10/15 |
| K1804 | BIG | 19.674 | -155.395 | *Vaccinium* | *reticulatum* | 01/10/15 |
| K1810 | BIG | 19.674 | -155.401 | *Vaccinium* | *reticulatum* | 01/10/15 |
| K1814 | BIG | 19.674 | -155.399 | *Vaccinium* | *reticulatum* | 01/10/15 |
| K1816 | BIG | 19.672 | -155.39 | *Vaccinium* | *reticulatum* | 01/10/15 |
| K1818 | BIG | 19.672 | -155.39 | *Vaccinium* | *reticulatum* | 01/10/15 |
| K1819 | BIG | 19.672 | -155.39 | *Vaccinium* | *reticulatum* | 01/10/15 |
| K1901 | BIG | 19.684 | -155.42 | *Vaccinium* | *reticulatum* | 01/10/15 |
| K1903 | BIG | 19.684 | -155.42 | *Vaccinium* | *reticulatum* | 01/10/15 |
| K1907 | BIG | 19.683 | -155.419 | *Vaccinium* | *reticulatum* | 01/10/15 |
| K1912 | BIG | 19.683 | -155.419 | *Vaccinium* | *reticulatum* | 01/10/15 |
| K2002 | BIG | 19.687 | -155.469 | *Vaccinium* | *reticulatum* | 01/10/15 |
| K2004 | BIG | 19.687 | -155.469 | *Vaccinium* | *reticulatum* | 01/10/15 |
| K2006 | BIG | 19.687 | -155.469 | *Vaccinium* | *reticulatum* | 01/10/15 |
| K2008 | BIG | 19.686 | -155.469 | *Vaccinium* | *reticulatum* | 01/10/15 |
| K2011 | BIG | 19.685 | -155.467 | *Vaccinium* | *reticulatum* | 01/10/15 |
| O102 | BIG | 19.619 | -155.359 | *Vaccinium* | *reticulatum* | 01/11/15 |
| O103 | BIG | 19.619 | -155.359 | *Vaccinium* | *reticulatum* | 01/11/15 |
| O106 | BIG | 19.619 | -155.359 | *Vaccinium* | *reticulatum* | 01/11/15 |
| O108 | BIG | 19.619 | -155.359 | *Vaccinium* | *reticulatum* | 01/11/15 |
| O110 | BIG | 19.62 | -155.359 | *Vaccinium* | *reticulatum* | 01/11/15 |
| O113 | BIG | 19.62 | -155.359 | *Vaccinium* | *reticulatum* | 01/11/15 |
| O117 | BIG | 19.62 | -155.359 | *Vaccinium* | *reticulatum* | 01/11/15 |
| O118 | BIG | 19.621 | -155.36 | *Vaccinium* | *reticulatum* | 01/11/15 |
| O119 | BIG | 19.622 | -155.36 | *Vaccinium* | *reticulatum* | 01/11/15 |
| O120 | BIG | 19.622 | -155.36 | *Vaccinium* | *reticulatum* | 01/11/15 |
| O201 | BIG | 19.63 | -155.362 | *Vaccinium* | *reticulatum* | 01/11/15 |
| O204 | BIG | 19.631 | -155.362 | *Vaccinium* | *reticulatum* | 01/11/15 |
| O207 | BIG | 19.631 | -155.362 | *Vaccinium* | *reticulatum* | 01/11/15 |
| O210 | BIG | 19.631 | -155.362 | *Vaccinium* | *reticulatum* | 01/11/15 |
| O212 | BIG | 19.631 | -155.362 | *Vaccinium* | *reticulatum* | 01/11/15 |
| O301 | BIG | 19.64 | -155.364 | *Vaccinium* | *reticulatum* | 01/11/15 |
| O302 | BIG | 19.64 | -155.364 | *Vaccinium* | *reticulatum* | 01/11/15 |
| O303 | BIG | 19.64 | -155.364 | *Vaccinium* | *reticulatum* | 01/11/15 |
| O310 | BIG | 19.64 | -155.364 | *Vaccinium* | *reticulatum* | 01/11/15 |

| | | | | | | |
|---|---|---|---|---|---|---|
| O313 | BIG | 19.641 | -155.364 | *Vaccinium* | *reticulatum* | 01/11/15 |
| O317 | BIG | 19.641 | -155.364 | *Vaccinium* | *reticulatum* | 01/11/15 |
| O401 | BIG | 19.649 | -155.366 | *Vaccinium* | *reticulatum* | 01/11/15 |
| O402 | BIG | 19.649 | -155.366 | *Vaccinium* | *reticulatum* | 01/11/15 |
| O404 | BIG | 19.649 | -155.366 | *Vaccinium* | *reticulatum* | 01/11/15 |
| O405 | BIG | 19.649 | -155.366 | *Vaccinium* | *reticulatum* | 01/11/15 |
| O406 | BIG | 19.649 | -155.366 | *Vaccinium* | *reticulatum* | 01/11/15 |
| O407 | BIG | 19.649 | -155.366 | *Vaccinium* | *reticulatum* | 01/11/15 |
| O409 | BIG | 19.65 | -155.366 | *Vaccinium* | *reticulatum* | 01/11/15 |
| O413 | BIG | 19.65 | -155.366 | *Vaccinium* | *reticulatum* | 01/11/15 |
| O415 | BIG | 19.65 | -155.366 | *Vaccinium* | *reticulatum* | 01/11/15 |
| O416 | BIG | 19.65 | -155.366 | *Vaccinium* | *reticulatum* | 01/11/15 |
| O419 | BIG | 19.65 | -155.366 | *Vaccinium* | *reticulatum* | 01/11/15 |
| O420 | BIG | 19.65 | -155.366 | *Vaccinium* | *reticulatum* | 01/11/15 |
| O424 | BIG | 19.65 | -155.366 | *Vaccinium* | *reticulatum* | 01/11/15 |
| O501 | BIG | 19.66 | -155.369 | *Vaccinium* | *reticulatum* | 01/11/15 |
| O504 | BIG | 19.66 | -155.369 | *Vaccinium* | *reticulatum* | 01/11/15 |
| O507 | BIG | 19.661 | -155.369 | *Vaccinium* | *reticulatum* | 01/11/15 |
| O510 | BIG | 19.661 | -155.369 | *Vaccinium* | *reticulatum* | 01/11/15 |
| O513 | BIG | 19.662 | -155.369 | *Vaccinium* | *reticulatum* | 01/11/15 |
| BI008 | FEF2 | 19.415 | -155.237 | *Vaccinium* | *sp.* | 07/21/14 |
| BI016 | FEF2 | 19.415 | -155.237 | *Vaccinium* | *sp.* | 07/21/14 |
| BI052 | FEF2 | 20.113 | -155.758 | *Vaccinium* | *calycinum* | 07/22/14 |
| BI097 | FEF2 | 19.676 | -155.329 | *Vaccinium* | *sp.* | 07/24/14 |
| BI098 | FEF2 | 19.675 | -155.329 | *Vaccinium* | *sp.* | 07/24/14 |
| BI106 | FEF2 | 19.672 | -155.338 | *Vaccinium* | *reticulatum* | 07/24/14 |
| BI119 | FEF2 | 19.676 | -155.384 | *Vaccinium* | *sp.* | 07/25/14 |
| BI123 | FEF2 | 19.673 | -155.385 | *Vaccinium* | *sp.* | 07/25/14 |
| BI134 | FEF2 | 19.649 | -155.372 | *Vaccinium* | *sp.* | 07/25/14 |
| BI142 | FEF2 | 19.647 | -155.369 | *Vaccinium* | *reticulatum* | 07/25/14 |
| BI153 | FEF2 | 20.09 | -155.737 | *Vaccinium* | *sp.* | 07/26/14 |
| K0031 | FEF2 | 22.15 | -159.642 | *Vaccinium* | *calycinum* | 06/15/15 |
| K0050 | FEF2 | 22.148 | -159.628 | *Vaccinium* | *calycinum* | 06/16/15 |
| K0061 | FEF2 | 22.154 | -159.618 | *Vaccinium* | *sp.* | 06/16/15 |
| K0069 | FEF2 | 22.147 | -159.618 | *Vaccinium* | *sp.* | 06/16/15 |
| Kan09 | FEF2 | 20.61 | -156.34 | *Vaccinium* | *sp.* | 06/10/15 |
| UW01 | FEF2 | 20.773 | -156.235 | *Vaccinium* | *reticulatum* | 11/06/14 |
| UW07 | FEF2 | 20.774 | -156.236 | *Vaccinium* | *calycinum* | 11/06/14 |
| wai16 | FEF2 | 20.799 | -156.253 | *Vaccinium* | *calycinum* | 11/04/14 |
| Mo006 | FEF3 | 21.119 | -156.93 | *Vaccinium* | *sp.* | 07/18/16 |
| Mo019 | FEF3 | 21.118 | -156.905 | *Vaccinium* | *sp.* | 07/18/16 |
| Mo029 | FEF3 | 21.119 | -156.901 | *Vaccinium* | *reticulatum* | 07/18/16 |
| Mo031 | FEF3 | 21.119 | -156.901 | *Vaccinium* | *sp.* | 07/18/16 |
| Mo070 | FEF3 | 21.102 | -156.906 | *Vaccinium* | *sp.* | 07/19/16 |
| Mo098 | FEF3 | 21.201 | -157.157 | *Vaccinium* | *sp.* | 07/20/16 |
| KAH4 | FEF3 | 21.556 | -157.869 | *Vigna* | *marina* | 07/29/16 |
| Mo024 | FEF3 | 21.118 | -156.905 | *Viola* | *sp.* | 07/18/16 |
| BI042 | FEF2 | 19.42 | -155.29 | *Wikstroemia* | *sp.* | 07/21/14 |
| BI088 | FEF2 | 19.448 | -154.862 | *Wikstroemia* | *sandwicensis* | 07/24/14 |
| BI209A | FEF2 | 19.615 | -155.927 | *Wikstroemia* | *sandwicensis* | 03/25/15 |
| BI209B | FEF2 | 19.615 | -155.927 | *Wikstroemia* | *sandwicensis* | 03/25/15 |
| BI234 | FEF2 | 19.615 | -155.931 | *Wikstroemia* | *sp.* | 03/25/15 |
| BI240 | FEF2 | 19.112 | -155.823 | *Wikstroemia* | *sp.* | 03/25/15 |
| K0001 | FEF2 | 22.034 | -159.669 | *Wikstroemia* | *furcata* | 06/15/15 |
| K0032 | FEF2 | 22.15 | -159.642 | *Wikstroemia* | *oahuensis* | 06/15/15 |
| HL9 | FEF3 | 21.324 | -157.742 | *Wikstroemia* | *oahuensis* | 07/21/16 |
| L1 | FEF3 | 21.635 | -157.943 | *Wikstroemia* | *oahuensis* | 07/02/16 |
| L7 | FEF3 | 21.596 | -157.958 | *Wikstroemia* | *oahuensis* | 07/02/16 |
| Mo060 | FEF3 | 21.1 | -156.914 | *Wikstroemia* | *oahuensis* | 07/19/16 |

| | | | | | | |
|---|---|---|---|---|---|---|
| POAR13 | FEF3 | 21.533 | -157.921 | *Wikstroemia* | *oahuensis* | 10/02/16 |
| Kan04A | FEF2 | 20.61 | -156.34 | *Nothocestrum* | *sp.* | 06/10/15 |
| wai26 | FEF2 | 20.801 | -156.253 | *Nothocestrum* | *latifolium* | 11/04/14 |
| K0041 | FEF2 | 22.149 | -159.645 | *Zanthoxylum* | *dipetalum* | 06/15/15 |

#### Table S3

PCoA results. “Geo” variables are PCNM vectors generated from the geographic distance matrix, and “Host” variables are PCNM vectors generated from the host phylogenetic distance matrix.

| **Variable** | $R^2$ | ***P*** |
|---|---|---|
| Julian date | 0.028 | 0.001 |
| NDVI | 0.066 | 0.001 |
| Evapotranspiration | 0.073 | 0.001 |
| Solar radiation | 0.041 | 0.001 |
| Elevation | 0.037 | 0.001 |
| Slope | 0.038 | 0.001 |
| Geo1 | 0.024 | 0.001 |
| Geo2 | 0.023 | 0.001 |
| Geo3 | 0.032 | 0.001 |
| Geo4 | 0.030 | 0.001 |
| Geo6 | 0.014 | 0.006 |
| Host3 | 0.106 | 0.001 |
| Host4 | 0.009 | 0.047 |
| Host5 | 0.020 | 0.001 |
| Host6 | 0.031 | 0.001 |

#### Table S4

Most common plant genera observed on each island. The four most common genera are shown, as well as their counts per island (in parenthesis).

| **Island** | **n Samp** | **Genus 1** | **Genus 2** | **Genus 3** | **Genus 4** |
|---|---|---|---|---|---|
| O’ahu | 123 | Metrosideros (9) | Coprosma (7) | Pittosporum (7) | Scaevola (6) |
| Hawai’i | 399 | Vaccimium (27) | Metrosideros (77) | Leptecophylla (73) | Coprosma (16) |
| Kaua’i | 80 | Scaevola (8) | Cyrtandra (5) | Metrosideros (4) | Vaccinium (4) |
| Maui | 51 | Acacia (3) | Coprosma (3) | Diospyros (3) | Dodonaea (3) |
| Moloka’i | 67 | Scaevola (9) | Vaccimium (4) | Dubautia (3) | Leptecophylla (3) |
